# Thermal-Acoustic Activation of Hydrophobic Polystyrene Supports for High-Efficiency Aqueous Solid-Phase Peptide Synthesis

**DOI:** 10.64898/2026.05.05.722603

**Authors:** Saranraj Krishnan, Aniruddha Kambekar, Janvi Khandelwal, Karthik Pushpavanam

## Abstract

Solid-phase peptide synthesis (SPPS) remains the dominant technique for peptide production. However, its reliance on hazardous organic solvents such as *N, N’*-dimethylformamide (DMF) and dichloromethane (DCM) results in an adverse environmental burden. One potential approach is replacing these organic solvents with water to reduce the hazardous solvent consumption and improve the environmental footprint of peptide production. This has led to the emergence of aqueous solid-phase peptide synthesis (ASPPS) approaches. Although successful, these approaches require specialized hydrophilic resins or modified building blocks, limiting their industrial applicability and scalability. Moreover, conventional hydrophobic polystyrene supports, remain the most widely used solid supports in industrial SPPS due to their high loading capacity, mechanical robustness, and low cost. These resins are generally considered incompatible with aqueous conditions. Here, we demonstrate that industrially relevant 2-chlorotrityl chloride (CTC) polystyrene resin can support efficient peptide coupling under fully aqueous conditions by integrating a precipitate-free 1-Ethyl-3-(3-dimethylaminopropyl) carbodiimide hydrochloride (EDC·HCl) and Oxyma activation system with a synergistic thermal–acoustic strategy. We posit that heating combined with ultrasonic irradiation likely promotes transient relaxation of the polystyrene matrix and enhances water penetration. This facilitates the diffusion of activated amino acid esters onto the hydrophobic resin required for coupling. The robustness of this aqueous methodology was validated through the synthesis of nine structurally diverse peptide sequences, including aromatic hydrogel-forming peptides, opioid peptides derived from enkephalins, toxin-inspired sequences, and a lipid-interacting fragment of *α*-synuclein. Analytical characterization by HPLC and MALDI-TOF mass spectrometry confirmed successful peptide assembly with high crude purity. We anticipate that this thermal–acoustic aqueous SPPS strategy provides a scalable and accessible pathway toward sustainable peptide manufacturing on classical hydrophobic supports with aqueous chemistry.

## Introduction

Peptides constitute a versatile class of biomolecules widely employed in therapeutics, diagnostics, catalysis, biomaterials, and molecular recognition.^1–3^ Peptides serve as critical biological mediators, acting as hormones, neurotransmitters, and growth factors that regulate numerous physiological processes.^3^ Their high specificity and relatively low toxicity make them attractive candidates for treating a wide range of conditions, including cancer, metabolic disorders such as diabetes and obesity, and rare diseases.^3^ The global peptide therapeutics market represents a rapidly expanding sector, valued at approximately $49.68 billion in 2026 and projected to exceed $70 billion by 2031.^4^ While blockbuster drugs such as glucagon-like peptide-1 (GLP-1) agonists (e.g., semaglutide) currently dominate market growth, the field is evolving toward innovative applications, including peptide–drug conjugates (PDCs) for targeted oncology and antimicrobial peptides (AMPs) to combat multidrug-resistant bacteria.^1,2^ Beyond medicine, peptides are increasingly utilized in the cosmetics industry for anti-aging skincare formulations and in the food industry for functional health benefits.^2^ Given the growing demand for peptide-based therapeutics and functional biomolecules, efficient and scalable synthetic methodologies are essential for their production.

Solid-phase peptide synthesis (SPPS), introduced by Merrifield, remains the dominant platform for peptide assembly due to its operational simplicity, compatibility with automation, and reproducibility. In SPPS, the peptide chain is constructed stepwise on an insoluble polymeric support, enabling iterative coupling and deprotection cycles.^5–7^ Despite its widespread use, conventional SPPS has a substantial environmental footprint because it relies heavily on large volumes of hazardous organic solvents, particularly *N, N’*-dimethylformamide (DMF) and dichloromethane (DCM). This accounts for the majority of chemical waste generated during peptide synthesis.^8^ Additionally, the European Union, as of December 12, 2023, has prohibited the manufacture, use, or sale of *N, N’*-dimethylformamide (DMF) unless exposure levels are restricted to 6 mg/m^3^.^9^ All these taken together have further intensified the need for greener peptide synthesis strategies.

In response to these concerns, significant efforts have been made to develop greener SPPS methodologies. Several strategies have focused on replacing DMF with less hazardous alternatives. *N*-Butylpyrrolidinone (NBP) has emerged as a promising drop-in replacement due to its low toxicity, strong solvating ability for protected amino acids, and compatibility with both polystyrene and PEG-based resins.^10–13^ *γ*-Valerolactone (GVL), a biomass-derived solvent obtained from cellulose, has also been reported as an effective medium for peptide coupling reactions.^11,12,14^ Similarly, Cyrene™ (dihydrolevoglucosenone), a renewable solvent produced from cellulose-derived feedstocks, offers high biodegradability and a favourable toxicity profile for amide bond formation.^11,12^ While these substitutions reduce the toxicity associated with conventional solvents, they still rely on organic solvent media and therefore do not fully address the environmental burden of SPPS. Other approaches have explored solvent-minimized or flow-based systems to reduce waste generation.^15^

More recently, Aqueous solid-phase peptide synthesis (ASPPS) has emerged as a promising alternative to replace bulk organic solvents with water. Early ASPPS strategies focused on modifying protecting groups or generating water-soluble Fmoc-amino acid derivatives to enhance solubility in aqueous media.^16–18^ While these studies confirm the feasibility of water-based peptide assembly, their broader adoption remains limited. Most reported methodologies rely on specialized hydrophilic supports which increase material costs and reduce accessibility for large-scale or resource-limited applications. In addition, hydrophilic resins typically undergo substantial swelling in aqueous conditions, which can complicate their use in packed-bed or continuous-flow peptide synthesis by altering bed volume and increasing pressure drop across the reactor.^19^

In contrast, hydrophobic polystyrene resins remain widely used solid supports in conventional SPPS due to their cross-linked polymer structure, high loading capacity, mechanical robustness, suppression of side reactions, compatibility with microwave-assisted synthesis, and economic advantages.^20,21^ However, these resins are generally considered incompatible with aqueous environments due to poor swelling, restricted reagent diffusion, and precipitation-driven pore blockage within the hydrophobic polymer matrix. Overcoming these limitations to enable efficient peptide synthesis on conventional hydrophobic polystyrene supports under aqueous conditions represents a critical challenge. Therefore, it is key to developing scalable and sustainable peptide synthesis strategies.

In this work, we demonstrate that conventional 2-chlorotrityl chloride (CTC) polystyrene resin can efficiently support fully aqueous peptide coupling when combined with a synergistic thermal–acoustic activation strategy. By integrating moderate heating and ultrasonic irradiation, we overcome hydrophobic resin collapse, enhance solvent penetration, and maintain homogeneous activation conditions in water. *N*-Methylmorpholine (NMM) is employed to generate water-compatible Fmoc-amino acid salts, while 1-Ethyl-3-(3-dimethylaminopropyl) carbodiimide hydrochloride (EDC·HCl) and Oxyma enables precipitate-free activation under aqueous conditions. By emphasizing water as the primary reaction medium and retaining inexpensive, industrially robust polystyrene supports,^22–24^ this strategy provides a scalable and economically viable pathway toward sustainable peptide manufacturing.

## Results and Discussion

To investigate whether efficient peptide coupling could be achieved under aqueous conditions on classical hydrophobic supports, we examined the performance of 2-chlorotrityl chloride (CTC) polystyrene resin using a thermal–acoustic activation strategy. This approach integrates moderate heating with ultrasonic irradiation to enhance reagent solubility and facilitate diffusion of activated amino acid intermediates into the resin matrix. In purely aqueous environments, CTC-polystyrene resin adopts a contracted, diffusion-restricted state due to insufficient solvation of its hydrophobic polymer backbone.^25^ Under ambient conditions, the polymer matrix remains poorly swollen, limiting reagent penetration into the internal pore structure and restricting mass transfer. We hypothesized that moderate heating (∼50 °C) could promote partial relaxation of the polymer chains at the resin surface, thereby improving accessibility of the internal reactive sites. When combined with ultrasonic irradiation (37–40 kHz, pulse mode), a synergistic effect emerges. Ultrasonic cavitation generates transient mechanical perturbations that likely disrupts the hydrophobic interchain interactions and increase the transient free volume within the polymer matrix.^26^ This dual activation mechanism effectively transforms the hydrophobic support into a more permeable matrix, facilitating diffusion of activated amino acid intermediates onto the resin under aqueous conditions.

The aqueous-based methodology for the synthesis of peptide sequences encompasses the processes of swelling, activation, coupling, deprotection, and washing steps conducted entirely in water. A key challenge in implementing this strategy lies in dissolving hydrophobic Fmoc-protected amino acids in aqueous media. To investigate this, we examined the dissolution behaviour of Fmoc-amino acids under different activation conditions (**Figure 1**). Initially, the Fmoc-amino acid was weighed and combined with the coupling reagent 1-ethyl-3-(3-dimethylaminopropyl) carbodiimide hydrochloride (EDC·HCl) and the additive Oxyma in a glass vial, where the reagents remained in the solid state prior to aqueous activation (**Figure 1A**). Manual mixing of the reagents with *N*-methylmorpholine (NMM) in water resulted in incomplete dissolution and visible undissolved particulates (**Figure 1B**). Similarly, only ultrasonic agitation at ambient temperature for 10 min did not significantly improve solubilization (**Figure 1C**). Heating the mixture to ∼50 °C with manual stirring led to partial dispersion, although undissolved material remained visible (**Figure 1D**). In contrast, simultaneous heating at ∼50 °C and ultrasonic irradiation produced a clear homogeneous yellow solution within 10 min (**Figure 1E**), indicating complete dissolution of the activated amino acid species.

**Figure 1.**
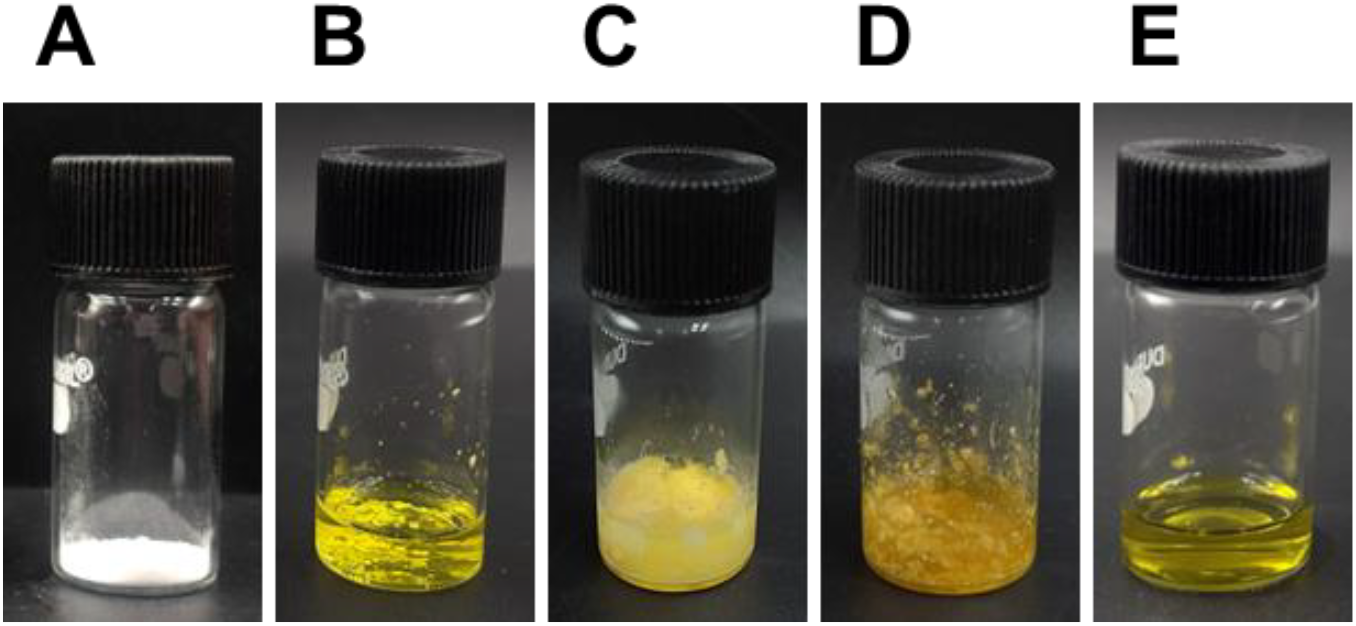
Dissolution behaviour of Fmoc-protected amino acid under various aqueous activation conditions. **(A)** Digital image of Fmoc-amino acid in the presence of 1-Ethyl-3-(3-dimethylaminopropyl) carbodiimide hydrochloride (EDC.HCl) and Oxyma prior to aqueous activation. **(B)** Fmoc-amino acid with EDC.HCl and Oxyma, N-methylmorpholine (NMM), in water mixed manually, showing incomplete dissolution and visible undissolved particulates. **(C)** The same mixture subjected to ultrasonic agitation at ambient temperature for 10 min, exhibiting persistent phase heterogeneity and incomplete solubilization. **(D)** Mixture heated to ∼50 °C with manual stirring, showing partial dispersion but continued presence of undissolved material. **(E)** Mixture heated to ∼50 °C and simultaneously subjected to ultrasonic irradiation for 10 min, resulting in a homogeneous yellow solution, indicating complete dissolution in water.

The proposed pathway for thermal–acoustic aqueous solid-phase peptide synthesis on hydrophobic supports (2-CTC resin) proceeds through six distinct stages (**Scheme 1**). The process begins by preparing a solution of NMM and water (**Scheme 1, Step-1**). To this mixture, Fmoc-amino acid is added to overcome its intrinsic insolubility in pure water (**Scheme 1, Step-2**). NMM acts as a base which generates a water-soluble carboxylate salt from the carboxylic acid group of Fmoc-amino acid through deprotonation. This process proceeds under heating (∼50 °C) and ultrasonic irradiation for 3–5 min. These conditions enhance dispersion and facilitate dissolution of the amino acid in the aqueous medium. Once the amino acid is fully dissolved, the coupling reagent EDC·HCl (a water-soluble carbodiimide) is introduced (**Scheme 1, Step-3**). EDC·HCl reacts with the carboxylate salt to generate an *O*-acylisourea intermediate. In this step, the EDC acts as a proton-transfer-activating agent, turning the amino acids oxygen into a good leaving group (LG). The use of the hydrochloride salt of EDC ensures adequate solubility and reactivity of the reagent in the aqueous medium. The highly reactive *O*-acylisourea intermediate is susceptible to hydrolysis in water. To prevent this side reaction, Oxyma is introduced as a stabilizing nucleophile (**Scheme 1, Step-4**). Oxyma reacts with the *O*-acylisourea intermediate to form a resonance-stabilized active ester. During this transformation, a water-soluble by-product, 1-ethyl-3-(3-dimethylaminopropyl)urea (EDU), is generated, which can be readily removed during subsequent washing steps. The above mixture containing the activated amino acid is introduced into the resin containing a growing peptide chain (**Scheme 1, Step 5**). The free amine on the peptide remains nucleophilic and attacks the carbonyl carbon of the active ester. This results in the formation of an amide bond. The reaction environment is maintained under mildly basic conditions by NMM. This formation of the amide bond extends the peptide chain by one amino acid residue (**Scheme 1, Step 6**). During this step, the Oxyma leaving group is released back into the solution. The resin is then sequentially washed to remove EDU and any residual reagents, leaving the hydrophobic support ready for the next deprotection and coupling step.

**Scheme 1.**
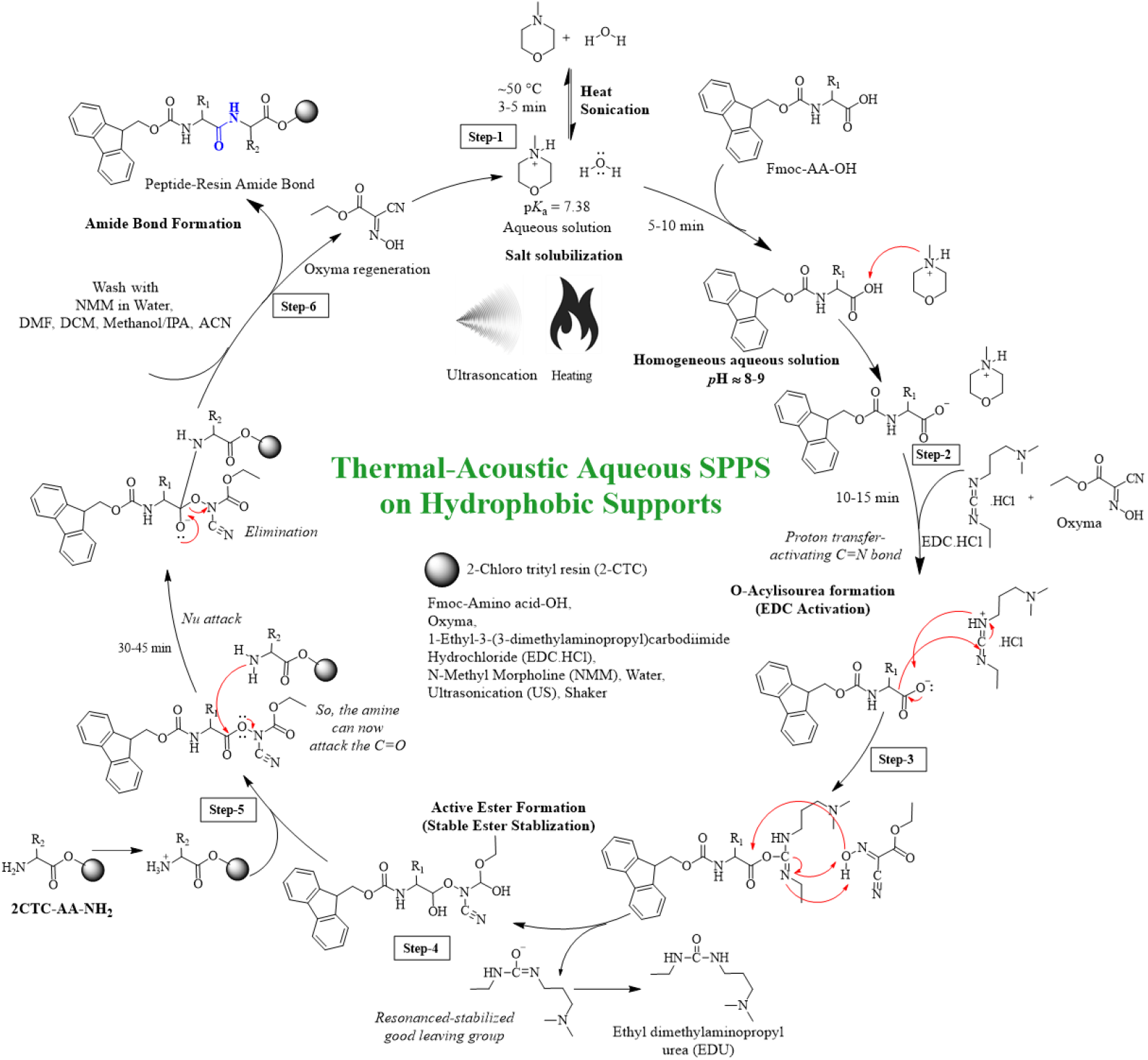
Mechanism of Thermal–Acoustic Aqueous Solid-Phase Peptide Synthesis on Polystyrene Support. Schematic representation of the coupling mechanism of Fmoc-protected amino acids on 2-chlorotrityl chloride (2-CTC) polystyrene resin under thermal–acoustic conditions. Initially, *N*-Methylmorpholine (NMM) is dissolved in water (**Step-1**), and the Fmoc-amino acid is further dissolved in this mixture under heating and ultrasonic irradiation, generating a homogeneous aqueous solution (**Step-2**). Upon addition of EDC·HCl, the carboxylate group is activated to form an *O*-acylisourea intermediate (**Step-3**), which is subsequently stabilized by Oxyma to produce a reactive active ester (**Step-4**). Under thermal–acoustic conditions, the activated ester diffuses into the hydrophobic resin matrix and reacts with the free amine (**Step-5**), resulting in amide bond formation and extension of the peptide chain (**Step-6**).

This methodology was validated using nine representative peptides in both Fmoc-protected and fully deprotected (free) forms (**Figure S1 & Table S1**). The study includes hydrogel-forming aromatic-rich peptide sequences YTPTY (Peptide-1) (**Figure 2A, 2C, S2 & S3**) and YTNTY (Peptide-2) (**Figure 2B, 2D, S4 & S5**). We have also synthesized opioid peptides YGGFL (Peptide-3) (**Figure S6 & S7**), YGGFLM (Peptide-4) (**Figure S8 & S9**), and its analogue YAGFLR (Peptide-5) (**Figure S10 & S11**) derived from Leu-enkephalin. Enkephalins are endogenous opioid neurotransmitters found in the brains of many animals, including humans. They primarily act as *δ*-opioid receptor agonists and play key roles in pain modulation, mood regulation, and stress response.^27^ Additionally, a model peptide IYGDKV (Peptide-6) (**Figure S12 & S13**) derived from Scorpion toxin II was prepared to represent bioactive toxin-inspired sequences, which are typically rich in charged residues and exhibit specific receptor interactions.^18,27^ Finally, a lipid-interacting segment DVFMKGLSKAK (Peptide-7) (**Figure S14 & S15**) derived from *Alpha*-synuclein (*α*S), was synthesized. α-Synuclein is a neuronal protein associated with membrane binding and is implicated in neurodegenerative disorders such as Parkinson’s disease. The chosen fragment is known for lipid-induced *α*-helix formation and membrane interaction.^28^ QRNA (Peptide-8) (**Figure S16 & S17**) and RGD (Peptide-9) (**Figure S18 & S19**) are arginine-containing peptide motifs commonly associated with cell adhesion to the extracellular matrix. The RGD motif is widely conserved and recognized across multiple species, ranging from Drosophila to humans.^29,30^ These sequences encompass sterically demanding, hydrophobic, charged, and amphiphilic residues, thereby rigorously evaluating diffusion, activation, and coupling efficiency under purely aqueous conditions.

**Figure 2.**
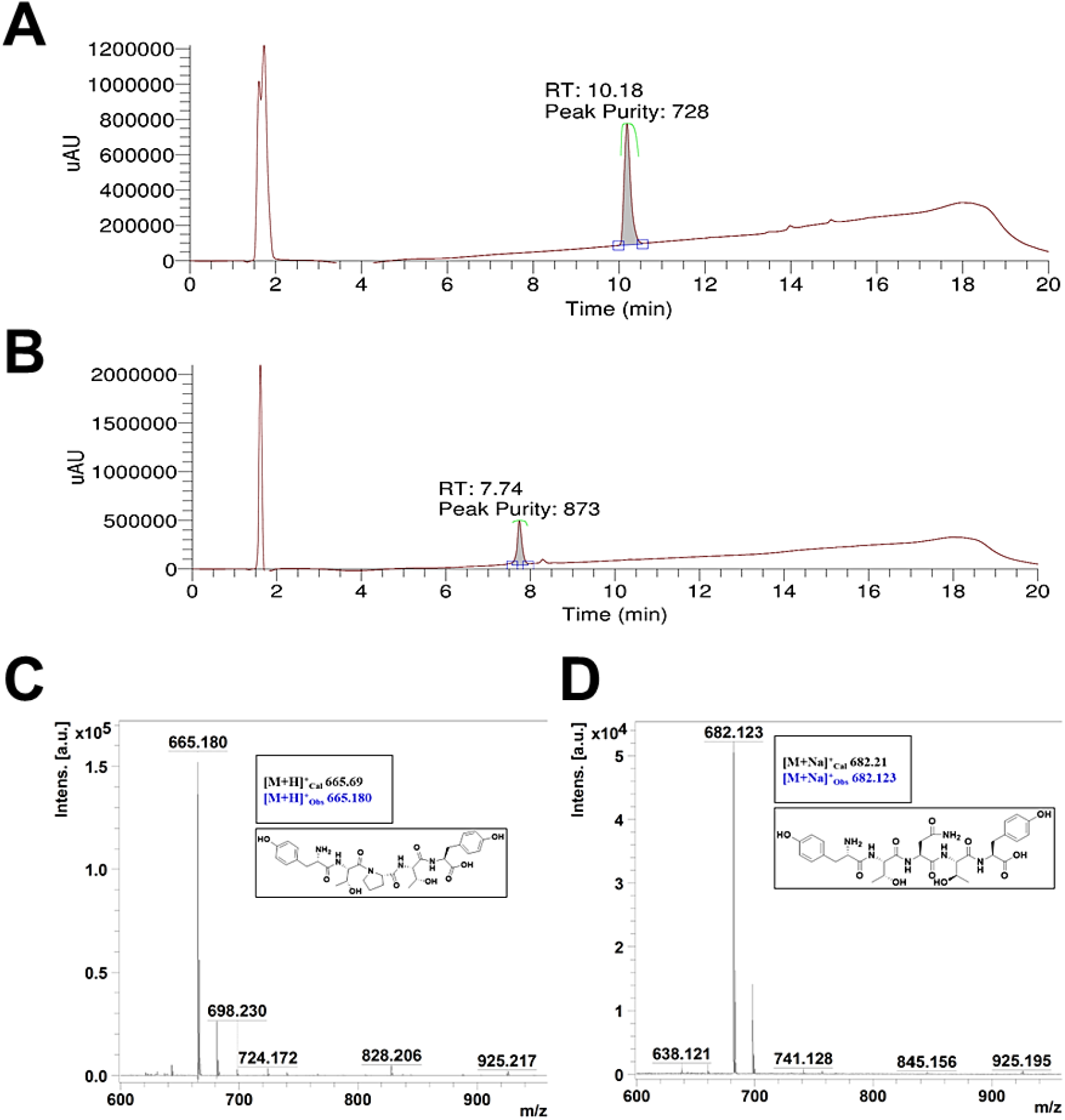
Analytical characterization of peptides synthesized by aqueous SPPS on 2-CTC polystyrene resin. (A, B) HPLC chromatograms of crude peptides: **(A)** Peptide-1 (YTPTY) and **(B)** Peptide-2 (YTNTY). Peptide samples were analyzed using a Thermo Fisher Scientific HPLC system equipped with a C_18_ column and UV detection at 220 nm. Separation was carried out using an acetonitrile–water mobile phase with a linear gradient of a binary solvent system (A: 95:5:0.1; B: 5:95:0.1) at a flow rate of 0.4 mL min^−1^ over a total run time of 20 min. (C, D) MALDI–TOF mass spectra confirming peptide identities for **(C)** Peptide-1 (YTPTY) and **(D)** Peptide-2 (YTNTY). The spectra display characteristic [M+H]^+^ and [M+Na]^+^ ion peaks, with experimentally observed masses in close agreement with the calculated molecular weights of the respective peptides.

High-performance liquid chromatography (HPLC) analysis of the crude peptides revealed well-defined major peaks with minimal side products, indicating satisfactory crude purity and absence of significant deletion sequences. The peptides exhibited distinct retention times (*t*_*R*_), with YTPTY (Peptide-1) eluting at 10.18 min (**Figure 2A**) and YTNTY (Peptide-2) at 7.74 min (**Figure 2B**). The solvent front/injection peak was consistently observed at 1.64 min and was well-resolved from the peptide signals. The difference in retention times can be attributed to the nature of the central amino acid residue in the peptide sequence. Peptide-1 contains a hydrophobic proline residue, which increases its interaction with the nonpolar C_18_ stationary phase, resulting in longer retention. In contrast, Peptide-2 contains the more polar asparagine residue, which exhibits weaker interactions with the hydrophobic stationary phase and therefore elutes earlier. Molecular integrity and composition were further verified by MALDI-TOF mass spectrometry, where the characteristic molecular ion peaks corresponding to [M+H]^+^, [M+Na]^+^, and [M+K]^+^ were observed for each peptide, matching the theoretical masses of the respective peptides (**Figure 2C, 2D & S2-19**). For the pentapeptide YTPTY (Peptide-1), the observed molecular weight of 665.18 Da closely matched the calculated value of 665.69 Da for the [M+H]^+^ (**Figure 2C**). Similarly, YTNTY (Peptide-2) exhibited an observed mass of 682.12 Da, consistent with the calculated [M+Na]^+^ ion at 682.21 Da (**Figure 2D**). Collectively, these analytical results confirm successful chain assembly and structural integrity under fully aqueous coupling conditions.

## Conclusion

This study demonstrates that conventional 2-chlorotrityl chloride (CTC) polystyrene resin can be effectively employed for peptide synthesis under fully aqueous coupling conditions by integrating a precipitate-free EDC·HCl and Oxyma activation system with a thermal–acoustic activation strategy. The combined application of moderate heating and ultrasonic irradiation mitigates hydrophobic contraction of the polystyrene matrix, enhances solvent penetration, and facilitates diffusion of activated amino acid intermediates onto the resin, enabling efficient amide bond formation on classical hydrophobic supports that are typically considered incompatible with aqueous environments. The robustness of this methodology was validated through the synthesis of nine structurally diverse peptides containing aromatic, hydrophobic, charged, and membrane-interacting residues. Analytical characterization by MALDI–TOF mass spectrometry and HPLC confirmed accurate chain assembly and satisfactory crude purity, demonstrating coupling efficiencies comparable to conventional DMF-based protocols. Importantly, replacing bulk organic solvents with water significantly reduces hazardous solvent consumption, waste generation, and overall environmental burden associated with SPPS. Beyond sustainability, the strategy retains the mechanical robustness, high loading capacity, and economic advantages of commercially available polystyrene supports, thereby preserving industrial practicality while improving environmental compatibility. We anticipate that this thermal–acoustic aqueous SPPS methodology expands the green chemistry toolkit for peptide synthesis and provides a scalable pathway toward more environmentally responsible peptide synthesis. Our future studies will focus on integration with automated synthesis platforms and evaluation of resin recyclability to further enhance process sustainability.

## Materials and Methods

### Materials

2-Chlorotrityl chloride resin (2-CTC, CAS Number: 934816-82-7), all the Fmoc-protected amino acids [Fmoc-Ala-OH (CAS Number: 35661-39-3), Fmoc-Cys(Trt)-OH (CAS Number: 103213-32-7), Fmoc-Asp(O*t*Bu)-OH (CAS Number: 71989-14-5), Fmoc-Glu(O*t*Bu)-OH (CAS Number: 71989-18-9), Fmoc-Phe-OH (CAS Number: 35661-40-6), Fmoc-Gly-OH (CAS Number: 29022-11-5), Fmoc-His(Trt)-OH (CAS Number: 109425-51-6), Fmoc-Ile-OH (CAS Number: 71989-23-6), Fmoc-Lys(Boc)-OH (CAS Number: 71989-26-9), Fmoc-Leu-OH (CAS Number: 35661-60-0), Fmoc-Met-OH (CAS Number: 71989-28-1), Fmoc-Asn(Trt)-OH (CAS Number: 132388-59-1), Fmoc-Pro-OH (CAS Number: 71989-31-6), Fmoc-Gln(Trt)-OH (CAS Number: 132327-80-1), Fmoc-Arg(Pbf)-OH (CAS Number: 154445-77-9), Fmoc-Ser(*t*Bu)-OH 71989-33-8), Fmoc-Thr(*t*Bu)-OH 71989-35-0), Fmoc-Val-OH 68858-20-8), Fmoc-Trp(Boc)-OH 143824-78-6), Fmoc-Tyr(*t*Bu)-OH 71989-38-3)], 1-ethyl-3-(3-dimethylaminopropyl)carbodiimide hydrochloride (EDC·HCl) (CAS Number: 25952-53-8), and ethyl cyanohydroxyiminoacetate (Oxyma) (CAS Number: 3849-21-6) were purchased from BLD Pharm. *N, N’*-diisopropylethylamine (DIPEA) (CAS Number: 387649), N-methylmorpholine (NMM) (CAS Number: 109-02-4), piperedine (PIP) (CAS Number: 110-89-4), (Acetic anhydride (CAS Number: 108-24-7) acetonitrile (ACN) (CAS Number: 75-05-8), methanol (MeOH) (CAS Number: 67-56-1), isopropyl alcohol (IPA) (CAS Number: 67-63-0), *N, N’*-dimethylformamide (DMF) (CAS Number: 68-12-2), dichloromethane (DCM) (CAS Number: 75-09-2), trifluoroacetic acid (TFA) (CAS Number: 76-05-1), triisopropylsilane (TIS) (CAS Number: T1533), diethyl ether (CAS Number: 60-29-7), and tert-butyl methyl ether (CAS Number: 1634-04-4) were obtained from Loba Chemie Private Limited. Milli-Q water was used as the reaction solvent unless otherwise specified. All reagents were used as received. Peptide synthesis was performed in polypropylene fritted syringe reactors under agitation using a Neuation shaker (iSHAK PS 10/20).

### Aqueous amino acid coupling protocol

Aqueous coupling reactions were performed using a thermal–acoustic activation strategy. *N*-Methylmorpholine (NMM) was dissolved in Milli-Q water and heated to ∼50 °C under ultrasonic irradiation until a clear solution was obtained. In a separate vial, the Fmoc-protected amino acid and Oxyma were combined, followed by addition of the preheated aqueous NMM solution. The mixture was sonicated until complete dissolution was achieved. EDC·HCl was subsequently added to generate the activated Oxyma ester in situ. The activated amino acid solution was immediately transferred to the resin bearing the free amine functionality. Coupling was allowed to proceed for 30-45 min at ∼50 °C under agitation. Following completion, the resin was washed sequentially with aqueous media. A 10% NMM in water wash was included to neutralize residual acidic species and preserve the integrity of the acid-labile CTC linker.

### Fmoc deprotection protocol

A 20% (v/v) aqueous piperidine solution was prepared and mixed until homogeneous. The resin was treated with this solution for two consecutive deprotection cycles of 15 min each, followed by the washing procedure.

### Peptide cleavage protocol

Peptides were cleaved, followed by two cycles, each cycle lasting 15 min, followed by a washing the resins using a TFA:TIS:water cleavage cocktail (95:2.5:2.5, v/v/v) for 1-2 h at room temperature. The cleavage mixture was filtered, and the filtrate was concentrated. Crude peptides were precipitated using cold diethyl ether, collected by centrifugation, and dried under vacuum.

### High-performance liquid chromatography (HPLC) Analysis

Peptide purity was analyzed using a Thermo Fisher Scientific Dionex UltiMate 3000 HPLC system coupled to an LCQ Fleet Ion Trap Mass Spectrometer. Separation was performed on a PepMap RSLC C_18_ analytical column (2 µm, 100 Å, 50 cm) with an Acclaim PepMap 100 precolumn (100 µm × 2 cm). The mobile phase consisted of solvent A (0.1% formic acid in Milli-Q water) and solvent B (80:20 acetonitrile:water containing 0.1% TFA). Peptides were eluted using a linear gradient from 0– 40% solvent B over 20 min at a flow rate of 0.4 mL min^-1^. Detection was performed at 220 nm. For direct analysis, 5 µL of the cleavage mixture was diluted 100-fold with water and acetonitrile, and 5 µL was injected. Data acquisition and processing were carried out using Xcalibur software (version 2.2 or later).

### MALDI-TOF Mass Spectrometry

Peptide samples were mixed with a matrix composed of sodium trifluoroacetate and 2,5-dihydroxybenzoic acid in tetrahydrofuran, drop-cast onto a MALDI target plate, and dried under ambient conditions. Mass spectra were recorded on a Bruker Autoflex MALDI-TOF instrument operated in linear positive mode over an m/z range of 300–3000. Data were processed using PolyTools software.

## AUTHOR CONTRIBUTIONS

The original idea, research concept, and experimental design were developed by S.K. S.K. and J.K. carried out experiments and data analysis. A.K. provided feedback and helped shape the analysis and the manuscript. K.P. supervised the overall research and provided critical feedback during manuscript writing. K.P. is thankful to the Department of Biotechnology-National Science Foundation RES/DBT/CL/P0340/2526/0018-4306 for the project funding.

## ACKNOWLEDGEMENTS

The authors would like to sincerely thank IIT Gandhinagar for providing the facilities and resources necessary to carry out this study. We thank CIF (Central Instrumentation Facility) for the instrumentation facilities.

## Notes

### Competing Interest Statement

The authors have declared no competing interest.

## REFERENCES

1. Xiao, W. et al. Advance in peptide-based drug development: delivery platforms, therapeutics and vaccines. Signal Transduct. Target. Ther. 10, 74 (2025).

2. Wang, M. et al. From precision synthesis to cross-industry applications: The future of emerging peptide technologies. Pharmacol. Res. 218, 107839 (2025).

3. Wang, L. et al. Therapeutic peptides: current applications and future directions. Signal Transduct. Target. Ther. 7, 48 (2022).

4. The Business Research Company. Peptide Therapeutics Market Report 2026. (2026).

5. Mitchell, A. R. Bruce Merrifield and solid-phase peptide synthesis: A historical assessment. Peptide Science 90, 175–184 (2008).

6. Merrifield, R. B. Solid-Phase Peptide Synthesis. III. An Improved Synthesis of Bradykinin ^*^. Biochemistry 3, 1385–1390 (1964).

7. Merrifield, R. B. Solid Phase Peptide Synthesis. I. The Synthesis of a Tetrapeptide. J. Am. Chem. Soc. 85, 2149–2154 (1963).

8. United Nations - Department of Economic and Social Affairs Sustainable Development. Transforming our world: the 2030 Agenda for Sustainable Development.

9. Sherwood, J., Albericio, F. & de la Torre, B. G. N, N -Dimethyl Formamide European Restriction Demands Solvent Substitution in Research and Development. ChemSusChem 17, (2024).

10. Malone, M. A., De Francisco Craparotta, M., Arnott, K. I. M., Jamieson, A. G. & Wade, N. Expanding the scope of sustainable peptide synthesis through post-linear synthesis reactions. Org. Biomol. Chem. 24, 1496–1502 (2026).

11. Scognamiglio, A. et al. Once Upon a Time Without DMF: Greener Paths in Peptide and Organic Synthesis. Molecules 31, 536 (2026).

12. Wegner, K., Barnes, D., Manzor, K., Jardine, A. & Moran, D. Evaluation of greener solvents for solid-phase peptide synthesis. Green Chem. Lett. Rev. 14, 153–164 (2021).

13. Lopez, J., Pletscher, S., Aemissegger, A., Bucher, C. & Gallou, F. N - Butylpyrrolidinone as Alternative Solvent for Solid-Phase Peptide Synthesis. Org. Process Res. Dev. 22, 494–503 (2018).

14. Kulkarni, C. et al. Optimized SPPS Process for Octreotide: Using Green Solvent GVL to Reduce Hazardous DMF and Ether Consumption. Org. Process Res. Dev. 29, 3126– 3137 (2025).

15. Ruhl, K. E. et al. Continuous-Flow Solid-Phase Peptide Synthesis to Enable Rapid, Multigram Deliveries of Peptides. Org. Process Res. Dev. 28, 2896–2905 (2024).

16. Hojo, K., Manabe, Y., Uda, T. & Tsuda, Y. Water-Based Solid-Phase Peptide Synthesis without Hydroxy Side Chain Protection. J. Org. Chem. 87, 11362–11368 (2022).

17. Phungula, A. et al. Aqueous Solid-Phase Peptide Synthesis (ASPPS) using Standard Fmoc/tBu-Protected Amino Acids. ACS Sustain. Chem. Eng. 13, 19833–19848 (2025).

18. Uth, C., Englert, S., Avrutina, O., Kolmar, H. & Knauer, S. Novel amino-Li resin for water-based solid-phase peptide synthesis. Journal of Peptide Science 29, (2023).

19. García-Martín, F. et al. ChemMatrix, a Poly(ethylene glycol)-Based Support for the Solid-Phase Synthesis of Complex Peptides. J. Comb. Chem. 8, 213–220 (2006).

20. García-Martín, F., Bayó-Puxan, N., Cruz, L. J., Bohling, J. C. & Albericio, F. Chlorotrityl Chloride (CTC) Resin as a Reusable Carboxyl Protecting Group. QSAR Comb. Sci. 26, 1027–1035 (2007).

21. Al Musaimi, O. et al. Towards green, scalable peptide synthesis: leveraging DEG-crosslinked polystyrene resins to overcome hydrophobicity challenges. RSC Adv. 14, 40255–40266 (2024).

22. Lopez, J. et al. Missing Link: Enabling Loading of 2-Chlorotrityl Chloride Resin in N - Butylpyrrolidinone as a Green Solvent. Org. Process Res. Dev. 26, 1450–1457 (2022).

23. Leko, M. et al. 2-Chlorotrityl Chloride and 4-Methylbenzhydryl Bromide Resin Loading Using the Mixture of Organic Solvents: A “Greener” Alternative to Dichloromethane and N, N -Dimethylformamide. Org. Process Res. Dev. 26, 144–148 (2022).

24. Alhassan, M., Al Musaimi, O., Collins, J. M., Albericio, F. & de la Torre, B. G. Cleaving protected peptides from 2-chlorotrityl chloride resin. Moving away from dichloromethane. Green Chemistry 22, 2840–2845 (2020).

25. Poly(Ethylene Glycol) Chemistry. (Springer US, Boston, MA, 1992). doi:10.1007/978-1-4899-0703-5.

26. Mottola, S. et al. Sustainable Ultrasound-Assisted Solid-Phase peptide synthesis (SUS-SPPS): Less Waste, more efficiency. Ultrason. Sonochem. 114, 107257 (2025).

27. Knauer, S. et al. Sustainable Peptide Synthesis Enabled by a Transient Protecting Group. Angew. Chem. Int. Ed. 59, 12984–12990 (2020).

28. Meade, R. M. et al. Stabilizing a Native Fold of Alpha-Synuclein with Short Helix-Constrained Peptides. JACS Au 5, 4321–4336 (2025).

29. Barczyk, M., Carracedo, S. & Gullberg, D. Integrins. Cell Tissue Res. 339, 269–280 (2010).

30. Davoodi, Z. & Shafiee, F. Internalizing RGD, a great motif for targeted peptide and protein delivery: a review article. Drug Deliv. Transl. Res. 12, 2261–2274 (2022).

