## Supplementary information data for "Thermal-Acoustic Activation of Hydrophobic Polystyrene Supports for High-Efficiency Aqueous Solid-Phase Peptide Synthesis": Supplementary Info - ASPPS.pdf

Karthik Pushpavanam, Ph.D.

Department of Chemical Engineering

Indian Institute of Technology Gandhinagar

Gujarat, India, 382055, India

### General Methods for Thermal–Acoustic Aqueous Solid-Phase Peptide Synthesis (ASPPS)

#### 1. General Synthetic Strategy

Peptides were synthesized using a manual-assisted peptide synthesis method on 2-chlorotrityl chloride-polystyrene (2-CTC-PS) resin using the conventional Fmoc protection strategy. The first amino acid was anchored to the resin via the reactive trityl chloride functionality. All coupling and deprotection cycles were performed under aqueous conditions using a Thermal–Acoustic protocol combining moderate heating (~50 °C) and ultrasonic irradiation (37–40 kHz, pulse mode).

Each synthetic cycle consisted of:

1. Resin swelling under thermal–acoustic conditions
2. Nα-Fmoc deprotection using 20% piperidine in water
3. Aqueous activation and coupling using EDC·HCl, Oxyma, and NMM
4. Sequential washing steps
5. Monitoring via Kaiser (ninhydrin) test and UV absorbance

Upon completion of chain assembly, peptides were cleaved using a TFA:TIS:water cocktail, precipitated, and lyophilized prior to characterization.

#### 2. Resin Swelling and Preconditioning

##### 2.1 Resin Weighing

The required amount of 2-CTC resin corresponding to a 0.1 mmol scale (resin loading: 0.3–4.0 mmol $g^{-1}$) was transferred into a reaction vial.

##### 2.2 Pre-Washing

To remove storage impurities, the resin was washed sequentially:

1. DMF (1×)
2. DCM (1×)
3. MeOH or IPA (1×)
4. MeCN (1×)

Each solvent was added to fully immerse the resin, the resin was agitated at 350 rpm for 1 min, and it was completely drained before proceeding.

##### 2.3 Preparation of 0.4 M NMM Aqueous Solution

*N*-Methylmorpholine (NMM) was dissolved in Milli-Q water to prepare a 0.4 M solution (2–3 equivalents to maintain pH 8–9). The solution was subjected to ultrasonic irradiation with heating (~50 °C) for ~10 min until a homogeneous solution was obtained.

#### 2.4 Thermal–Acoustic Swelling

The warm NMM solution (~50 °C) was added to the pre-washed resin. Swelling was conducted for 30–45 min under mild sonication at ~50 °C followed by shaking.

Thermal input enhances polymer chain mobility, while ultrasonic cavitation induces transient mechanical perturbation, increasing transient free volume within the hydrophobic matrix.

### 3. Fmoc Deprotection (Aqueous Conditions)

#### 3.1 Preparation of Deprotection Solution

A 20% (v/v) piperidine solution in water was prepared and mixed thoroughly until homogeneous.

#### 3.2 Deprotection Cycles

Two consecutive deprotection cycles were performed:

1. Add sufficient 20% piperidine solution (~2 ml) to immerse resin
2. Subject to mild heating (~50 °C) and sonication
3. Allow reaction for 15 min/cycle

#### 3.3 Post-Deprotection Washing

After deprotection, the resin was washed sequentially:

1. DMF (1×)
2. DCM (1×)
3. MeOH/IPA (1×)
4. MeCN (1×)

Complete drainage after each wash.

### 4. Preparation of Activated Amino Acid Solution (Thermal–Acoustic Pre-Activation)

Fmoc-protected amino acid (3 equiv.) and Oxyma (4 equiv.) were combined in a separate vial. Freshly prepared warm 0.4 M NMM aqueous solution was added.

The mixture was subjected to ultrasonic irradiation (37–40 kHz) at ~50 °C for 3–10 min until a completely homogeneous solution was obtained.

Under these conditions:

1. The carboxylic acid is deprotonated to form a water-soluble carboxylate salt.
2. pH was maintained between 8–9.
3. Hydrophobic amino acids remain dispersed without aggregation.
4. Bulky residues such as Tyr(*t*Bu) and Thr(*t*Bu) remain fully dissolved.

Thermal input stabilizes ionic species and prevents precipitation ("crashing out") of activated intermediates.

[NOTE: Frequency: A standard frequency of 37-40 kHz is generally used for SPPS]. It provides enough power to penetrate the mixture and/or resin without being so aggressive (like lower 20 kHz frequencies) that it crushes the mixture and/or beads. Mode: Use pulse mode (e.g., 4 seconds ON / 2 seconds OFF) to prevent the solution from overheating, which could lead to racemization or resin degradation]

**5. Carbodiimide Activation and Active Ester Formation**

EDC·HCl (3–4 equiv.) was added to the amino acid and Oxyma mixture under continued sonication at ~50 °C.

Mechanistic Sequence:

1. EDC reacts with the carboxylate salt to form a high-energy *O*-acylisourea intermediate.
2. Oxyma rapidly converts this intermediate into a stabilized active ester.
3. The water-soluble urea byproduct EDU is released.

Importantly, EDU contains a tertiary amine that remains protonated under mildly basic 0.4 M NMM conditions, rendering it highly water-soluble and preventing precipitation within the resin matrix. This maintains a homogeneous reaction phase and avoids pore blockage commonly observed with DIC-derived DIU in aqueous systems. The formation of a clear activated solution indicated successful in situ generation of the active ester.

**6. Acoustic-Assisted Coupling to Resin**

The activated amino acid solution was immediately transferred to the resin bearing the free amine (Fmoc-deprotected).

Coupling conditions: Shaking at 350 rpm, 30-45 min reaction time, pH maintained at 8–9.

Mechanistic Considerations: Under acoustic irradiation, Cavitation induces mechanical perturbation of the polystyrene matrix, the transient free volume increases, and the activated esters diffuse onto hydrophobic resin. The free amine nucleophilically attacks the active ester carbonyl, forming the amide bond and regenerating Oxyma.

[NOTE: pH 8-9: This range is high enough to ensure the *N*-terminal amine of your growing peptide is deprotonated (active) so it can attack the incoming amino acid, but low enough to minimize the risk of base-catalysed side reactions]

**7. Post-Coupling Basic Wash**

A 10% NMM in water wash was incorporated after coupling.

CTC resin is highly acid-labile; residual acidic species from EDC·HCl or Oxyma may promote premature cleavage. The basic wash neutralizes residual acidic components, preserving linker integrity and minimizing product loss.

**8. Washing Procedure After Each Coupling**

After coupling, the resin is filtered and washed sequentially:

1. DMF (1×)
2. DCM (1×)
3. MeOH/IPA (1×)
4. MeCN (1×)

Each solvent was added to immerse the resin, agitated for 1–2 min, and completely drained before proceeding.

The resin was dried under vacuum or nitrogen flow before the next synthetic cycle.

**9. Monitoring of Peptide Elongation**

Each coupling and deprotection step was monitored using:

1. Kaiser (ninhydrin) test, the absence of blue coloration confirms the consumption of the free amine and successful chain elongation.
2. UV absorbance analysis

Blue coloration indicated presence of free primary amine; absence of colour change confirmed successful amine consumption and chain elongation.

**10. Cleavage of Peptide from Resin**

Following final deprotection, the resin was transferred to a syringe and dried under $N_2$.

Cleavage cocktail composition:

- 95% TFA
- 2.5% TIS
- 2.5% Milli-Q water

Procedure:

1. Add 5 mL cleavage cocktail.
2. Agitate for 1 h.
3. Precipitate in cold *tert*-butyl methyl ether.
4. Repeat cleavage with additional 5 mL cocktail.
5. Refrigerate precipitate overnight.
6. Centrifuge at 6000 rpm (HERMLE Z 36 HK) for 1 h.
7. Wash pellet with excess *tert*-butyl methyl ether.

**11. Lyophilization**

Milli Q water was added to the peptide precipitate. The sample was deep-frozen in liquid nitrogen and lyophilized using a LABOGENE ScanVac CoolSafe freeze dryer. Lyophilized peptide powders were stored at 4 °C until further analysis.

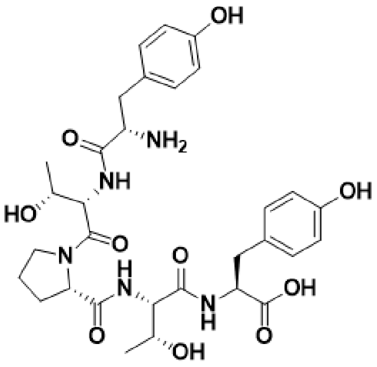

H-Tyr-Thr-Pro-Thr-Tyr-OH
(Peptide-1)

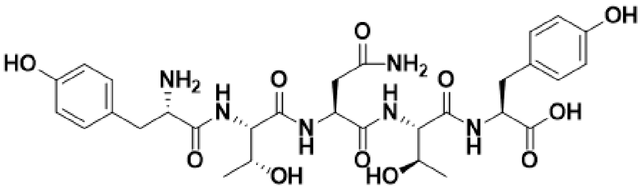

H-Tyr-Thr-Asn-Thr-Tyr-OH
(Peptide-2)

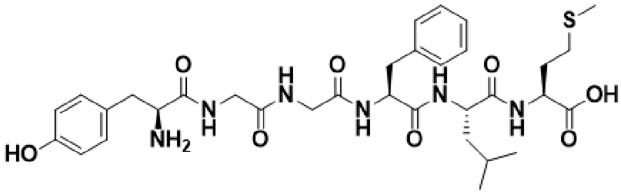

H-Tyr-Gly-Gly-Phe-Met-OH
(Peptide-4)

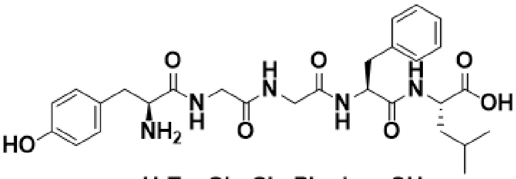

H-Tyr-Gly-Gly-Phe-Leu-OH
(Peptide-3)

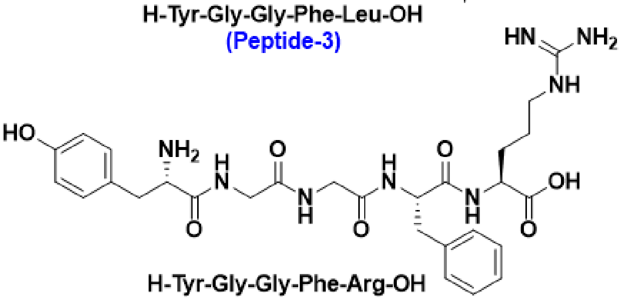

H-Tyr-Gly-Gly-Phe-Arg-OH
(Peptide-5)

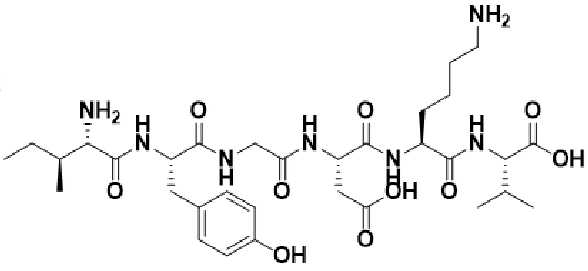

H-Ile-Tyr-Gly-Asp-Lys-Val-OH
(Peptide-6)

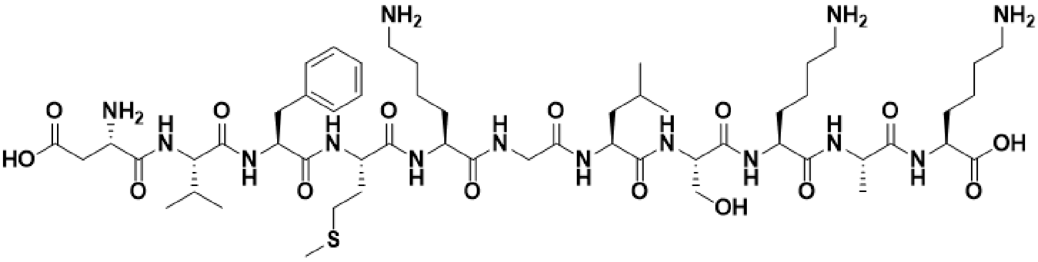

H-Asp-Val-Phe-Met-Lys-Gly-Leu-Ser-Lys-Ala-Lys-OH
(Peptide-7)

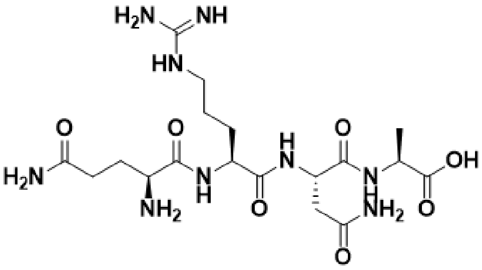

H-Gln-Arg-Asn-Ala-OH
(Peptide-8)

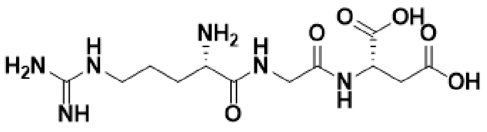

H-Arg-Gly-Asp-OH
(Peptide-9)

**Figure S1. Structures of synthesized peptides.** Chemical structures of the peptides used in this study: Peptide-1 (YTPTY), Peptide-2 (YTNTY), Peptide-3 (YGGFL), Peptide-4 (YGGFLM), Peptide-5 (YAGFLR), Peptide-6 (IYGDKV), Peptide-7 (DVFMKGLSKAK), Peptide-8 (QRNA), and Peptide-9 (RGD).

**Table S1.** Characteristics analysis of peptide sequences. The peptides termed Peptide-XF and Peptide-X represent Fmoc-containing peptides and the other without, respectively. X stands for the identifier number.

| Peptide code | Peptide sequence | Functionality | Cal. MW | Obs. MW |
|---|---|---|---|---|
| Peptide-1F | Fmoc-Tyr-Thr-Pro-Thr-Tyr-OH | Hydrogel | 887.78 | 887.356 |
| Peptide-1 | H-Tyr-Thr-Pro-Thr-Tyr-OH | | 665.69 | 665.180 |
| Peptide-2F | Fmoc-Tyr-Thr-Asn-Thr-Tyr-OH | Hydrogel | 904.52 | 904.425 |
| Peptide-2 | H-Tyr-Thr-Asn-Thr-Tyr-OH | | 682.21 | 682.123 |
| Peptide-3F | Fmoc-Tyr-Gly-Gly-Phe-Leu-OH | Leu-enkephalin | 800.87 | 800.584 |
| Peptide-3 | H-Tyr-Gly-Gly-Phe-Leu-OH | | 578.27 | 578.165 |
| Peptide-4F | Fmoc-Tyr-Gly-Gly-Phe-Leu-Met-OH | Modified Met-enkephalin | 947.245 | 947.38 |
| Peptide-4 | H-Tyr-Gly-Gly-Phe-Leu-Met-OH | | 687.31 | 687.495 |
| Peptide-5F | Fmoc-Tyr-Ala-Gly-Phe-Leu-Arg-OH | Opioid receptor | 948.45 | 948.230 |
| Peptide-5 | H-Tyr-Ala-Gly-Phe-Leu-Arg-OH | | 726.39. | 726.203 |
| Peptide-6F | Fmoc-Ile-Tyr-Gly-Asp-Lys-Val-OH | Scorpion toxin II | 948.44 | 948.249 |
| Peptide-6 | H-Ile-Tyr-Gly-Asp-Lys-Val-OH | | 694.37 | 694.231 |
| Peptide-7F | Fmoc-Asp-Val-Phe-Met-Lys-Gly-Leu-Ser-Lys-Ala-Lys-OH | Lipid binding region | 1445.74 | 1445.500 |
| Peptide-7 | H-Asp-Val-Phe-Met-Lys-Gly-Leu-Ser-Lys-Ala-Lys-OH | | 1223.67 | 1223.496 |
| Peptide-8F | Fmoc-Gln-Arg-Asn-Ala-OH | Cell adhesion | 710.76 | 710.237 |
| Peptide-8 | H-Gln-Arg-Asn-Ala-OH | | 488.25 | 488.640 |
| Peptide-9F | Fmoc-Arg-Gly-Asp-OH | Cell adhesion | 569.23 | 569.534 |
| Peptide-9 | H-Arg-Gly-Asp-OH | | 347.16 | 347.434 |

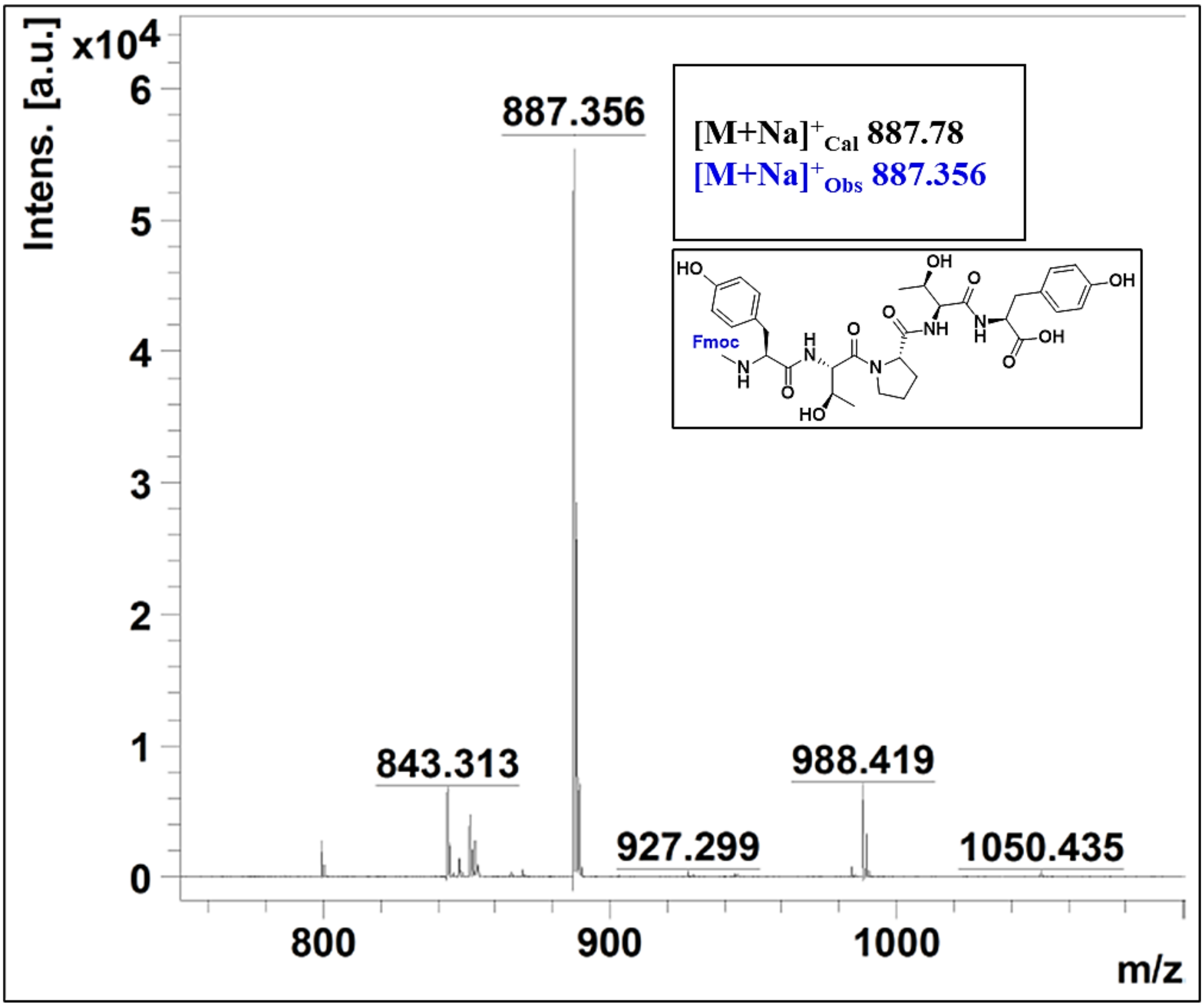


**Figure S2.** MALDI spectrum of Fmoc-Tyr-Thr-Pro-Thr-Tyr-OH in a mixture of acetonitrile: water, expected molecular weight is 887.78, and the observed molecular weight *via* MALDI is 887.356.

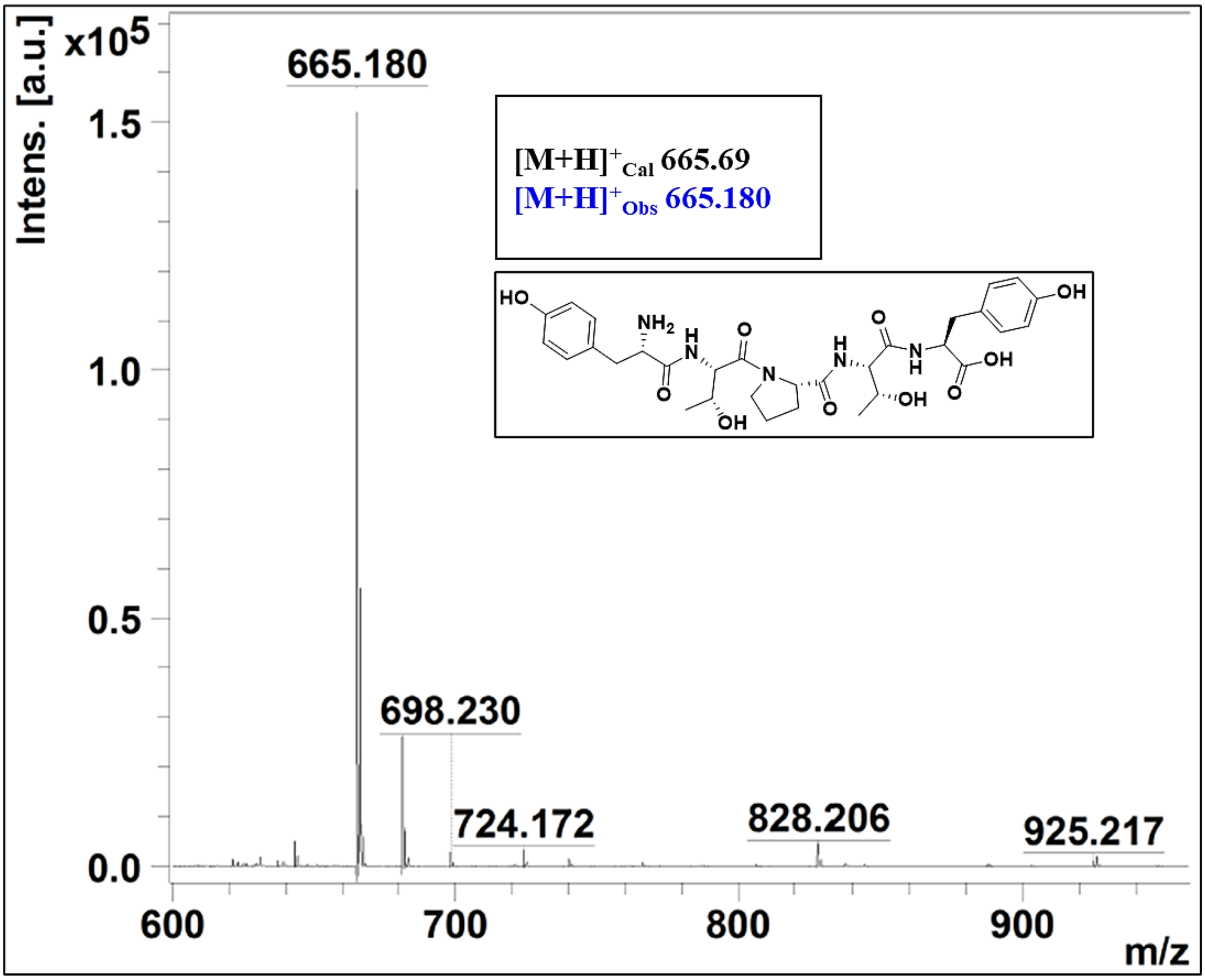


**Figure S3.** MALDI spectrum of H-Tyr-Thr-Pro-Thr-Tyr-OH in a mixture of acetonitrile: water, expected molecular weight is 665.69, and the observed molecular weight *via* MALDI is 665.180.

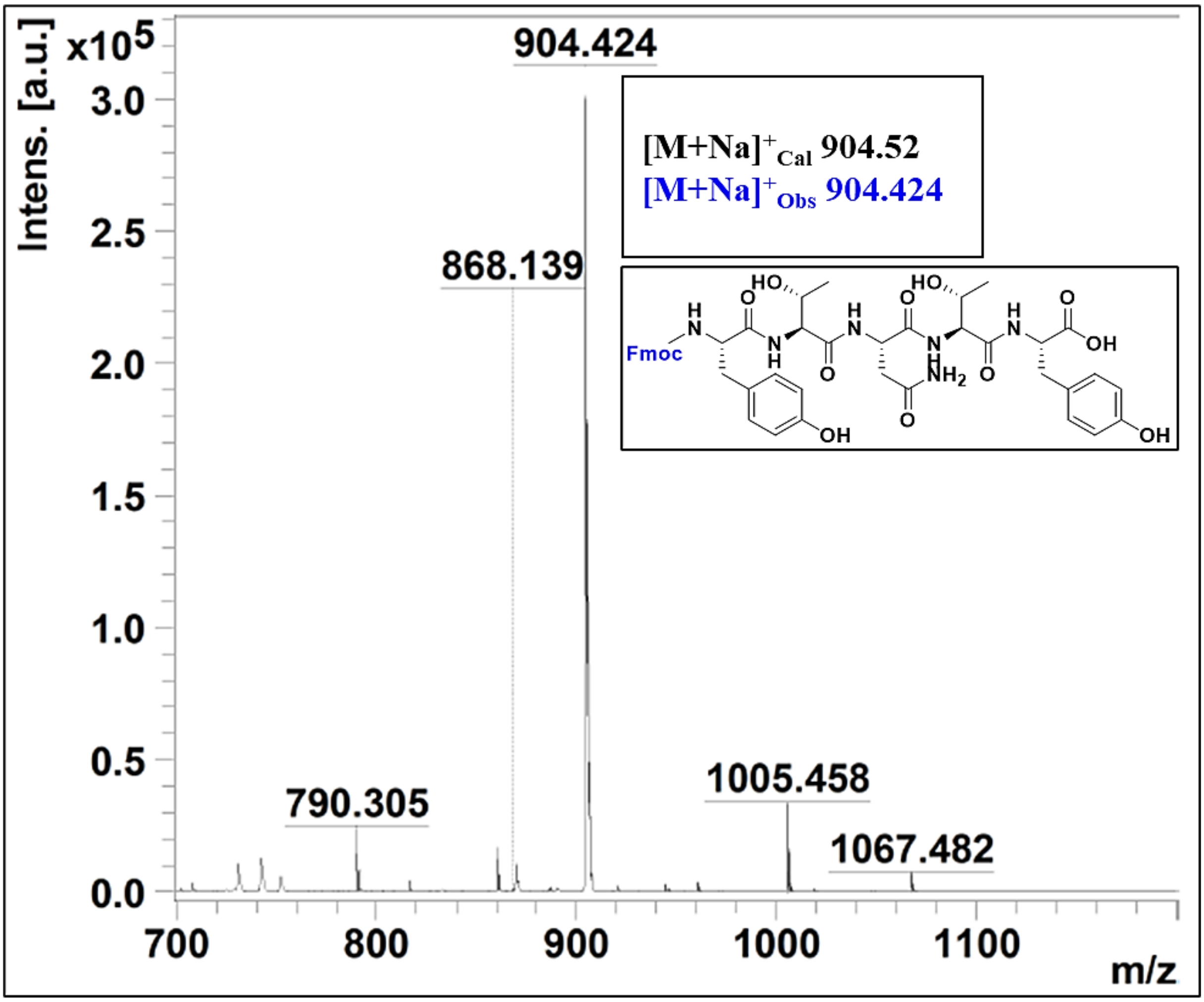


**Figure S4.** MALDI spectrum of Fmoc-Tyr-Thr-Asn-Thr-Tyr-OH in a mixture of acetonitrile: water, expected molecular weight is 904.52, and the observed molecular weight *via* MALDI is 904.424.

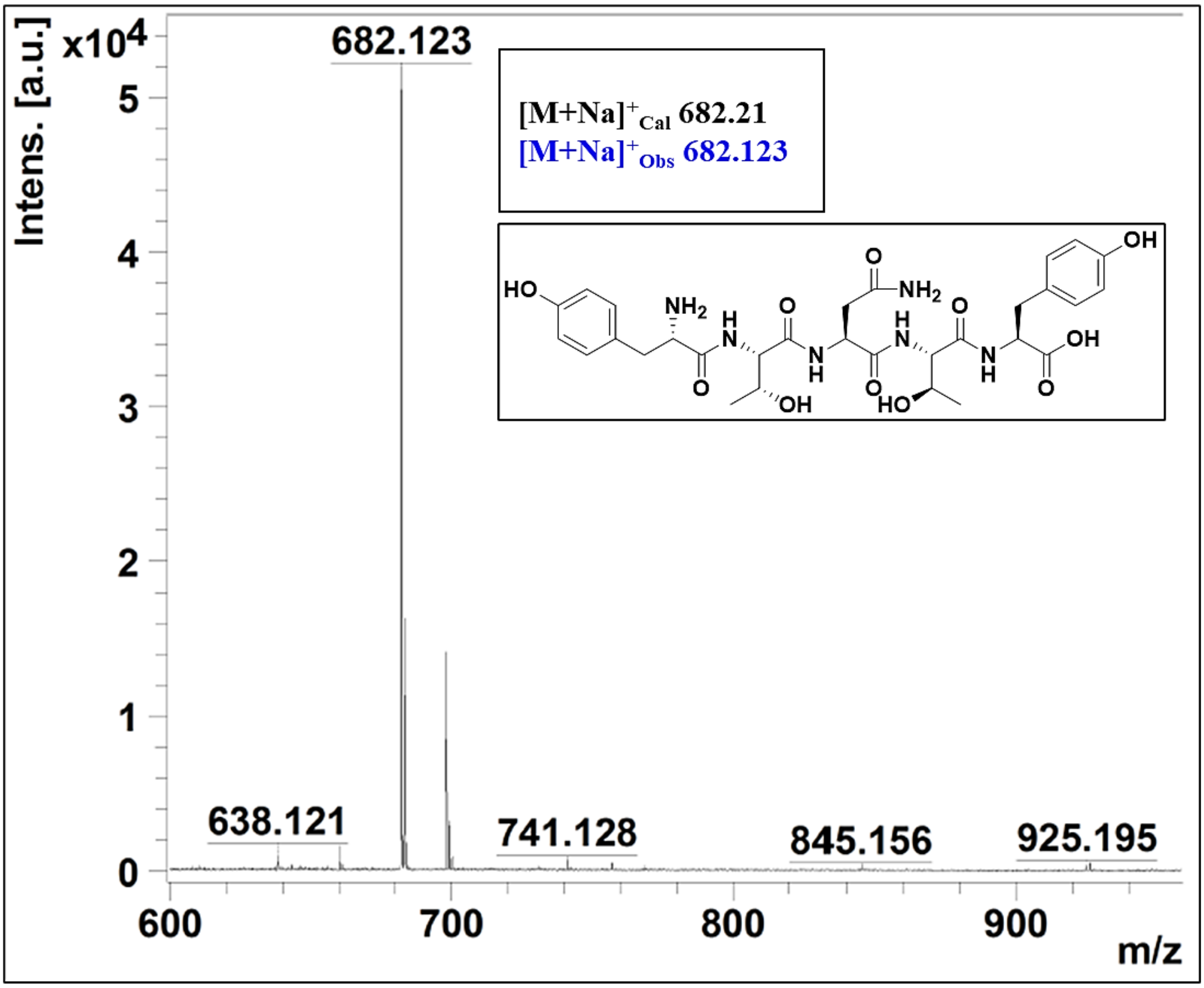


**Figure S5.** MALDI spectrum of H-Tyr-Thr-Asn-Thr-Tyr-OH in a mixture of acetonitrile: water, expected molecular weight is 682.21, and the observed molecular weight *via* MALDI is 682.123.

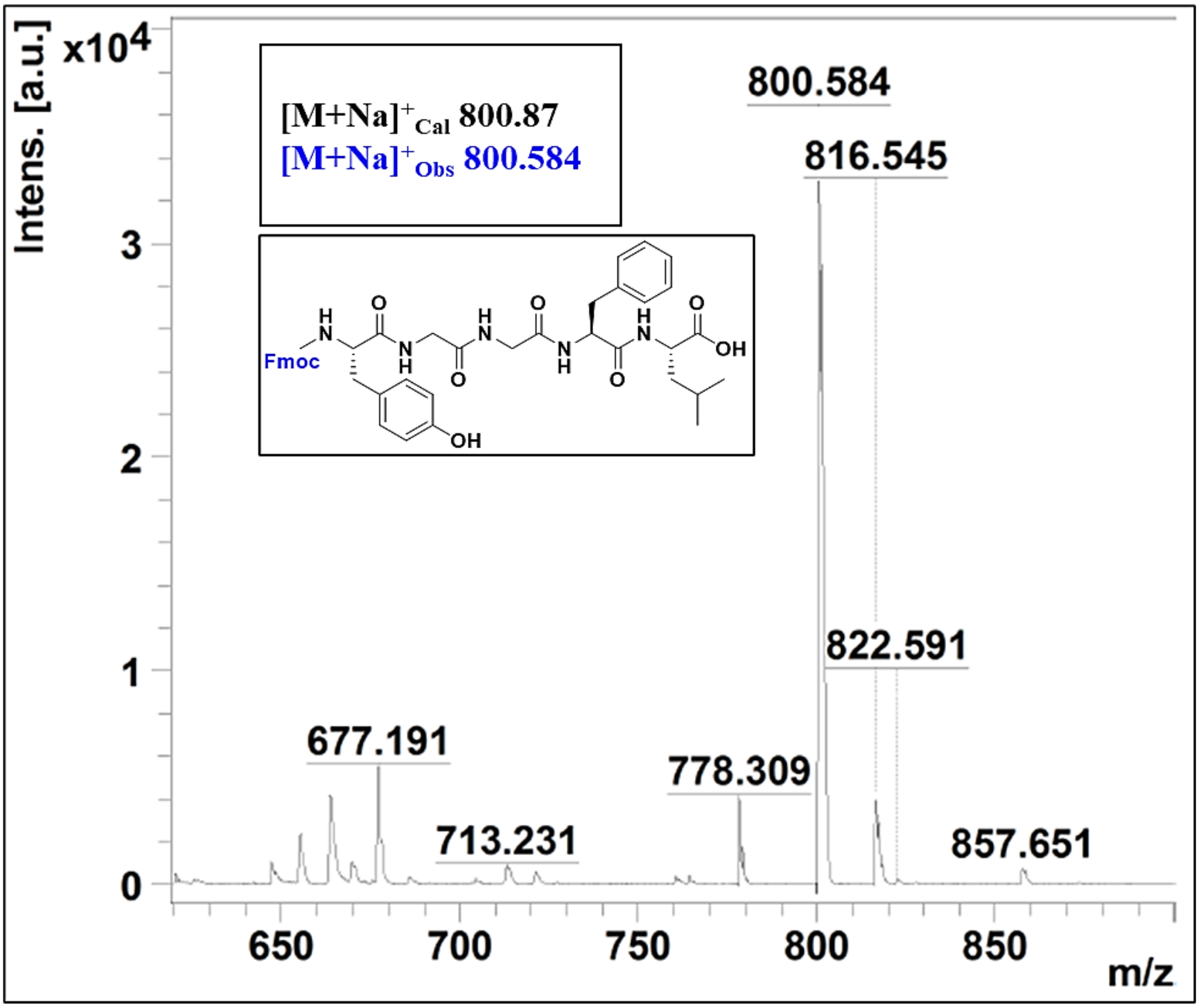


**Figure S6.** MALDI spectrum of Fmoc-Tyr-Gly-Gly-Phe-Leu-OH in a mixture of acetonitrile: water, expected molecular weight is 800.87, and the observed molecular weight *via* MALDI is 800.584.

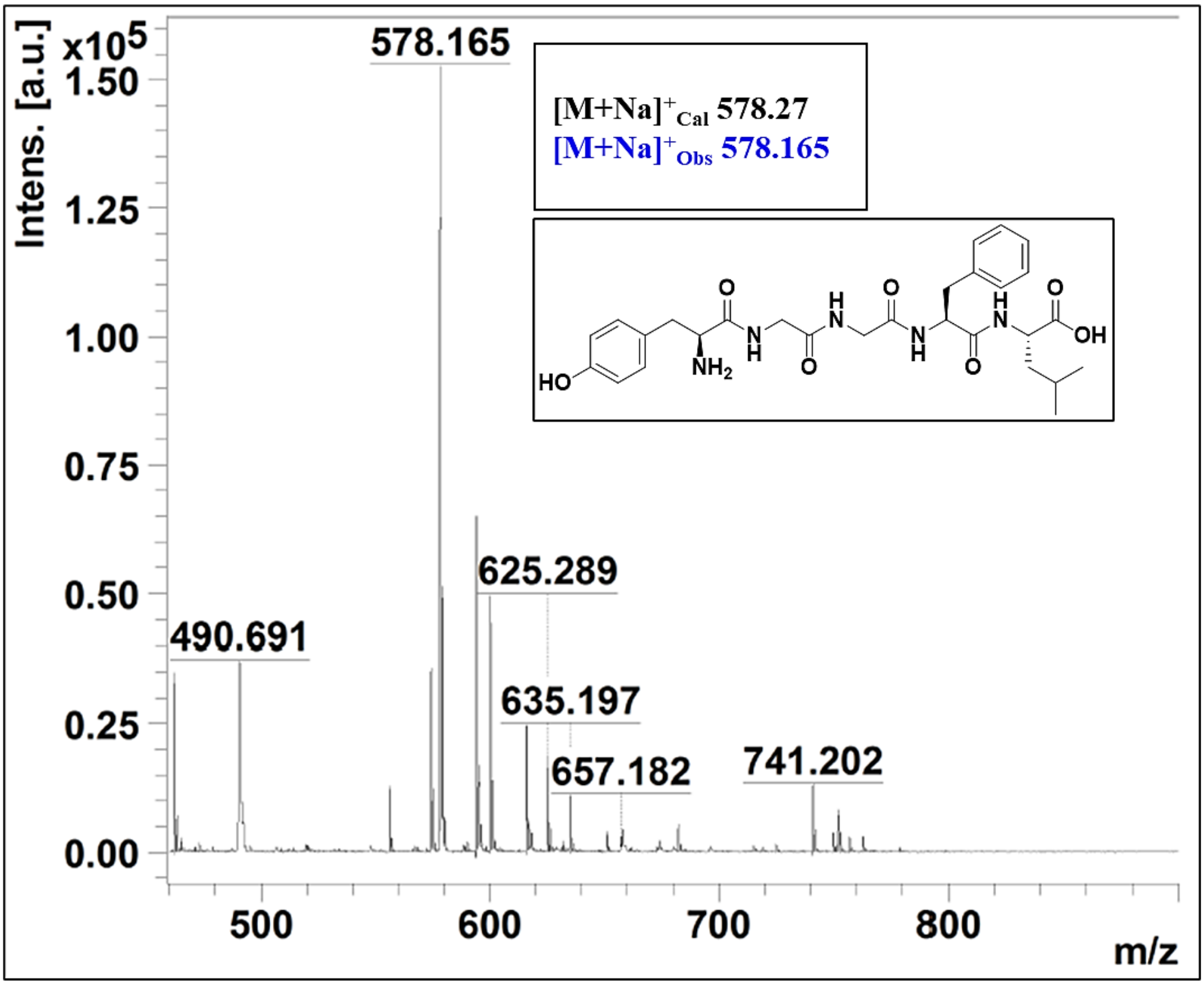


**Figure S7.** MALDI spectrum of H-Tyr-Gly-Gly-Phe-Leu-OH in a mixture of acetonitrile: water, expected molecular weight is 578.27, and the observed molecular weight *via* MALDI is 578.165.

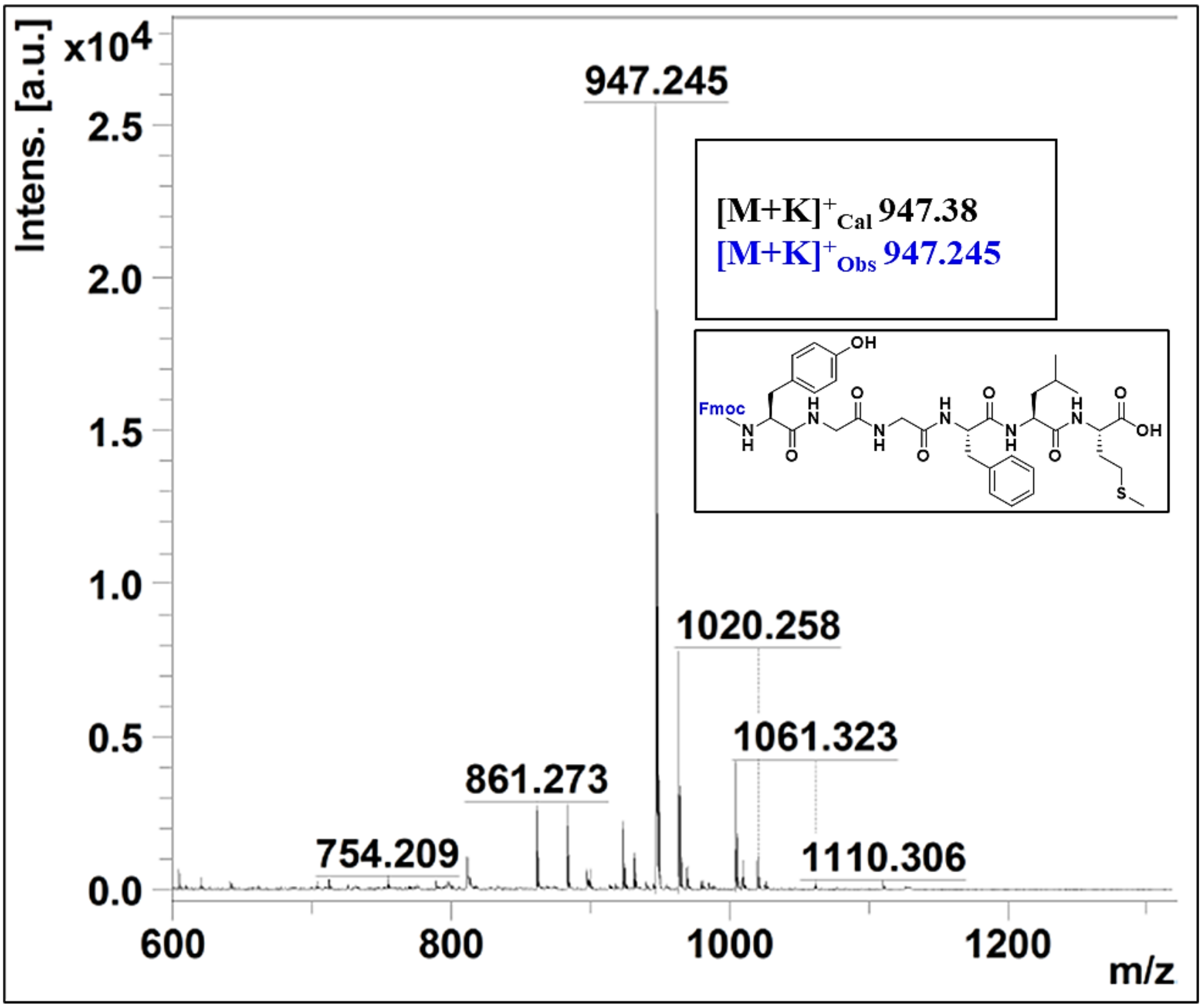


**Figure S8.** MALDI spectrum of Fmoc-Tyr-Gly-Gly-Phe-Leu-Met-OH in a mixture of acetonitrile: water, expected molecular weight is 947.38, and the observed molecular weight *via* MALDI is 947.245.

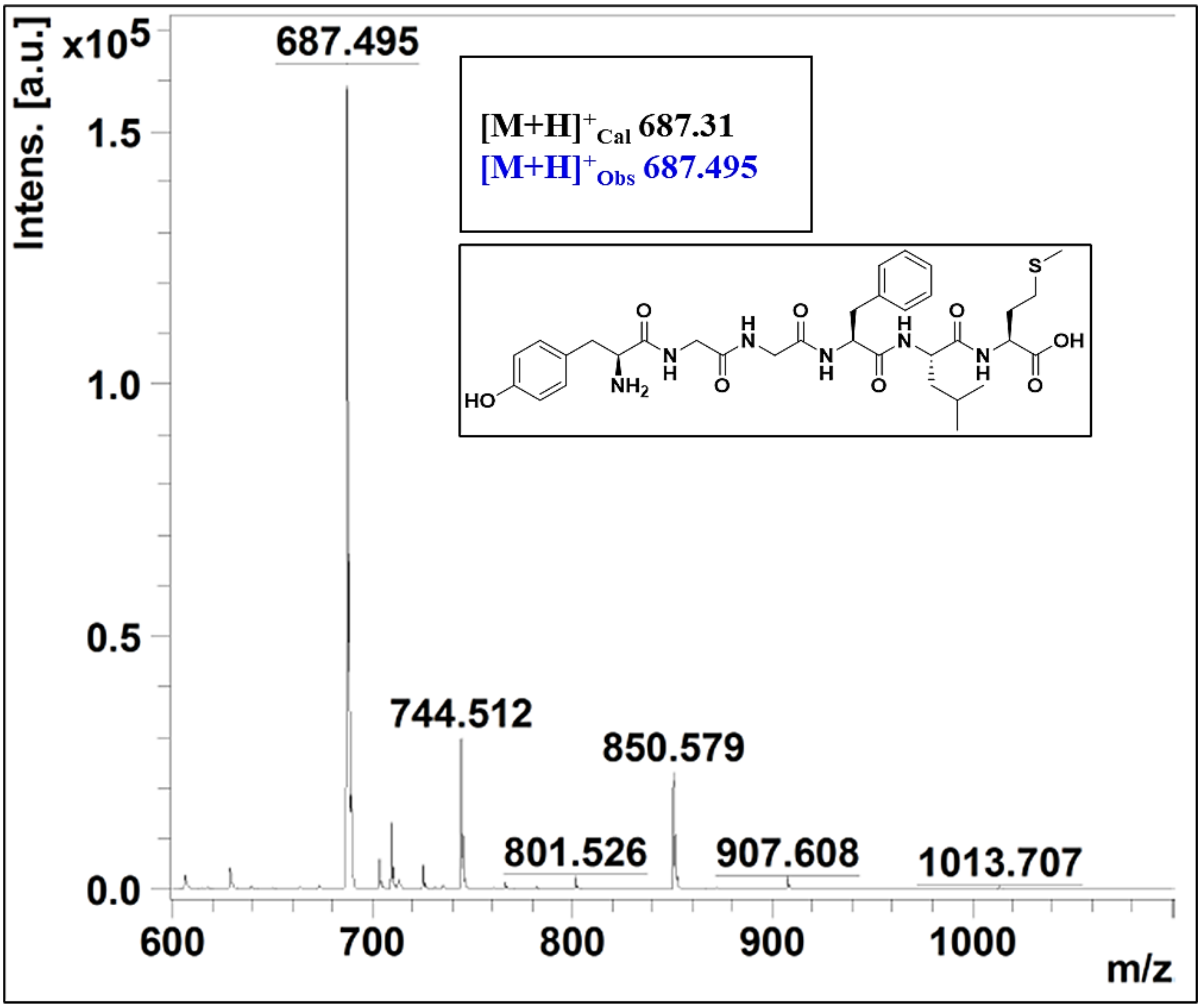


**Figure S9.** MALDI spectrum of H-Tyr-Gly-Gly-Phe-Leu-Met-OH in a mixture of acetonitrile: water, expected molecular weight is 687.31, and the observed molecular weight *via* MALDI is 687.495.

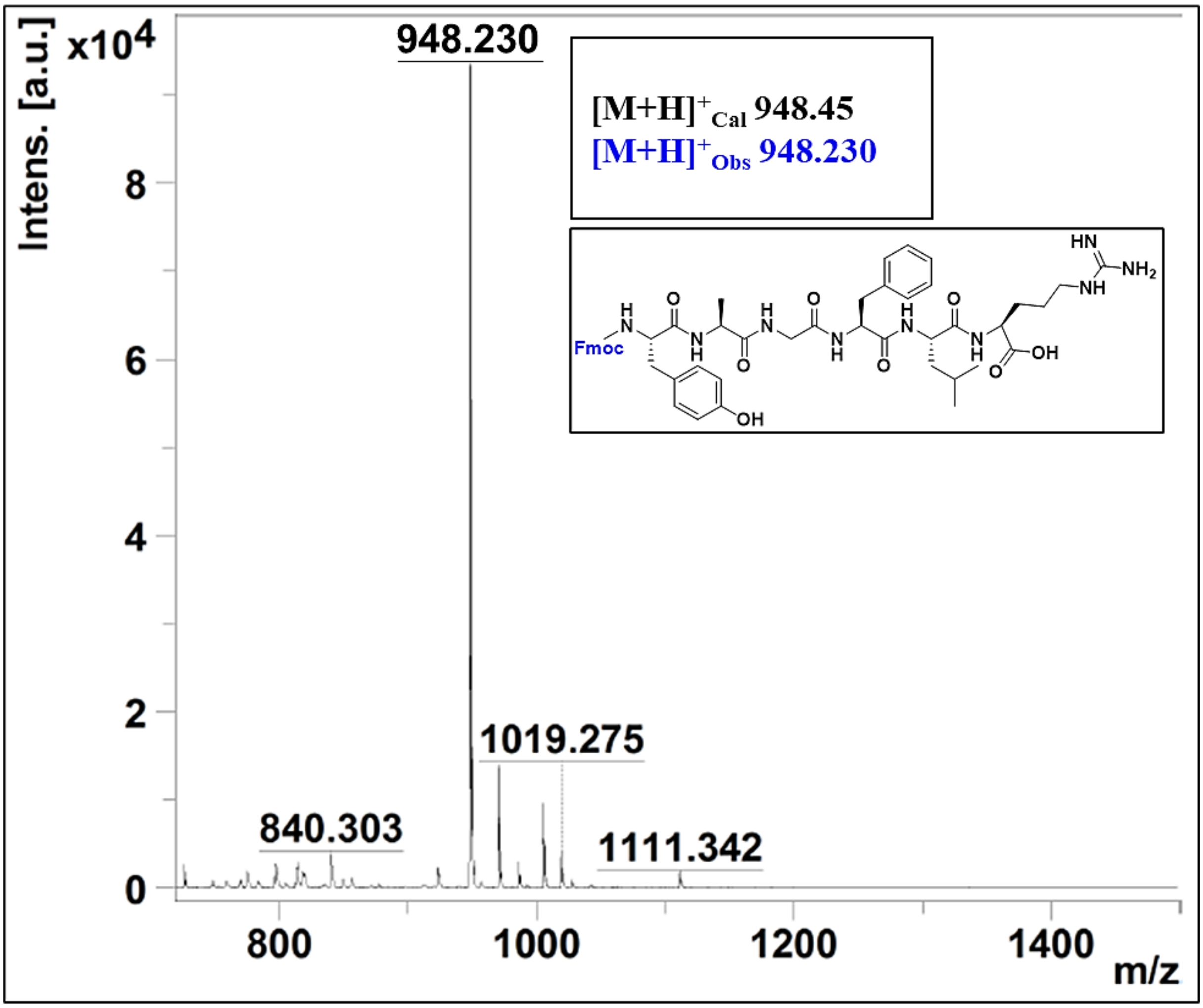


**Figure S10.** MALDI spectrum of Fmoc-Tyr-Ala-Gly-Phe-Leu-Arg-OH in a mixture of acetonitrile: water**,** expected molecular weight is 948.45, and the observed molecular weight *via* MALDI is 948.230.

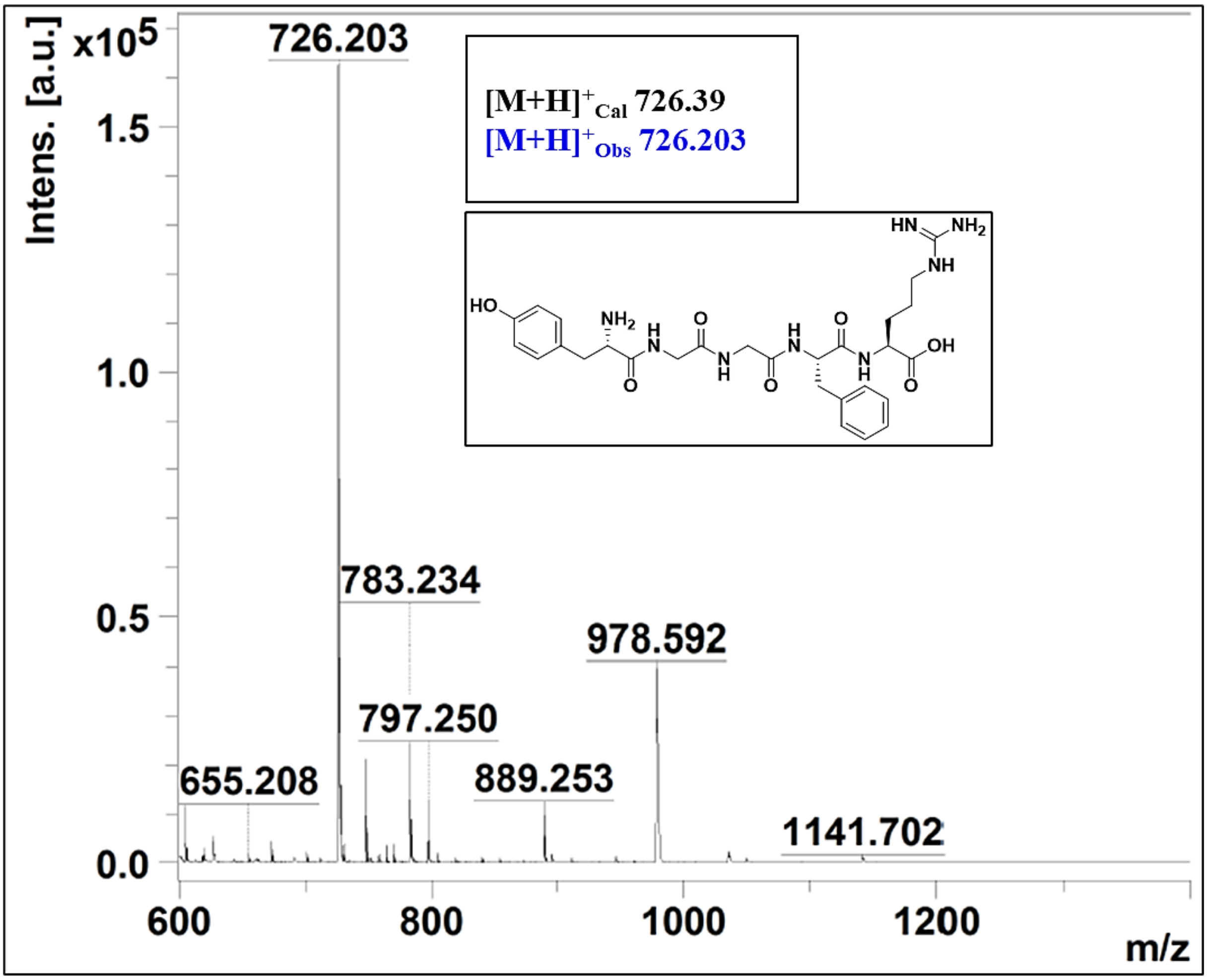


**Figure S11.** MALDI spectrum of H-Tyr-Ala-Gly-Phe-Leu-Arg-OH in a mixture of acetonitrile: water, expected molecular weight is 726.39, and the observed molecular weight *via* MALDI is 726.203.

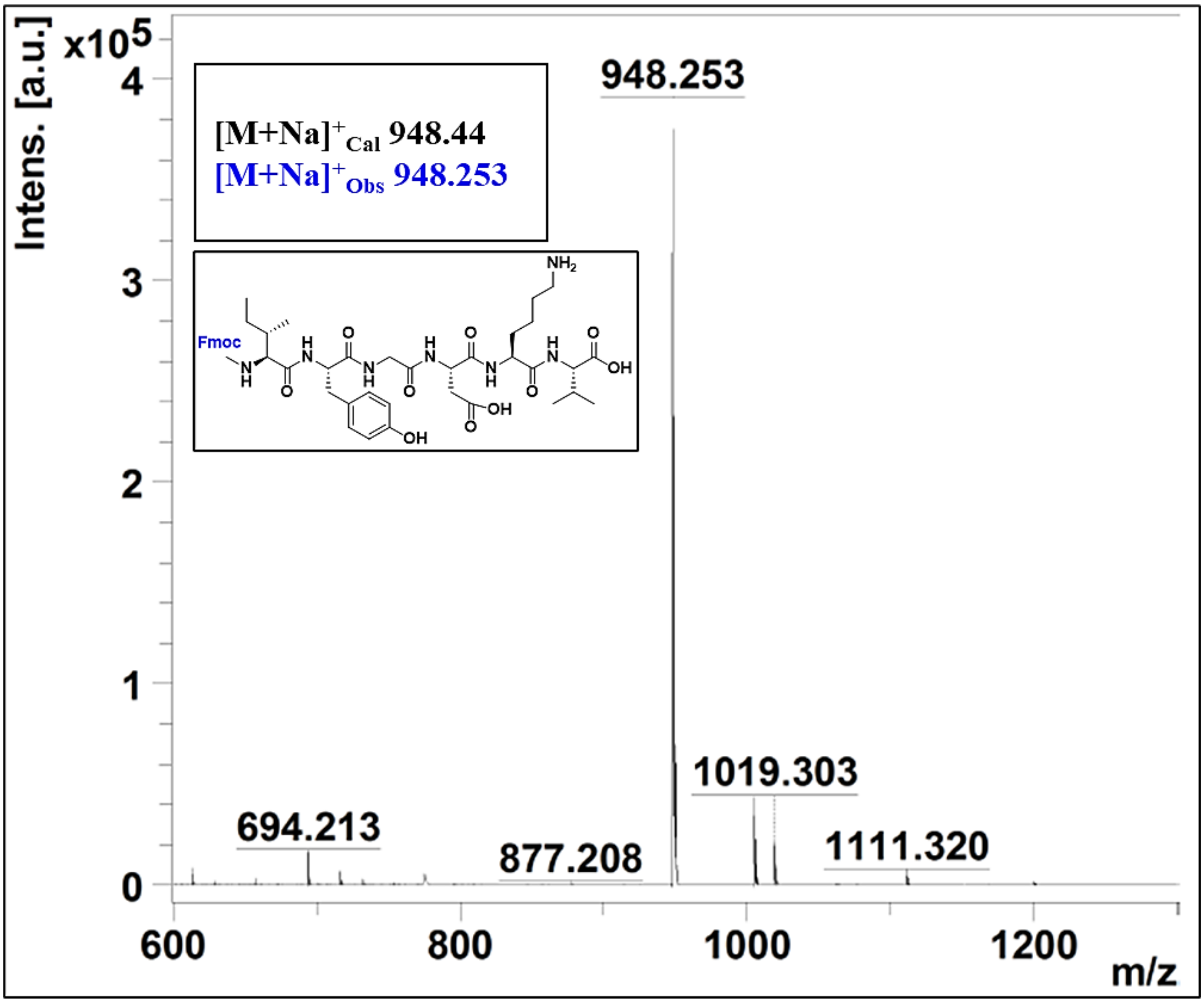


**Figure S12.** MALDI spectrum of Fmoc-Ile-Tyr-Gly-Asp-Lys-Val-OH in a mixture of acetonitrile: water, expected molecular weight is 948.44, and the observed molecular weight *via* MALDI is 948.253.

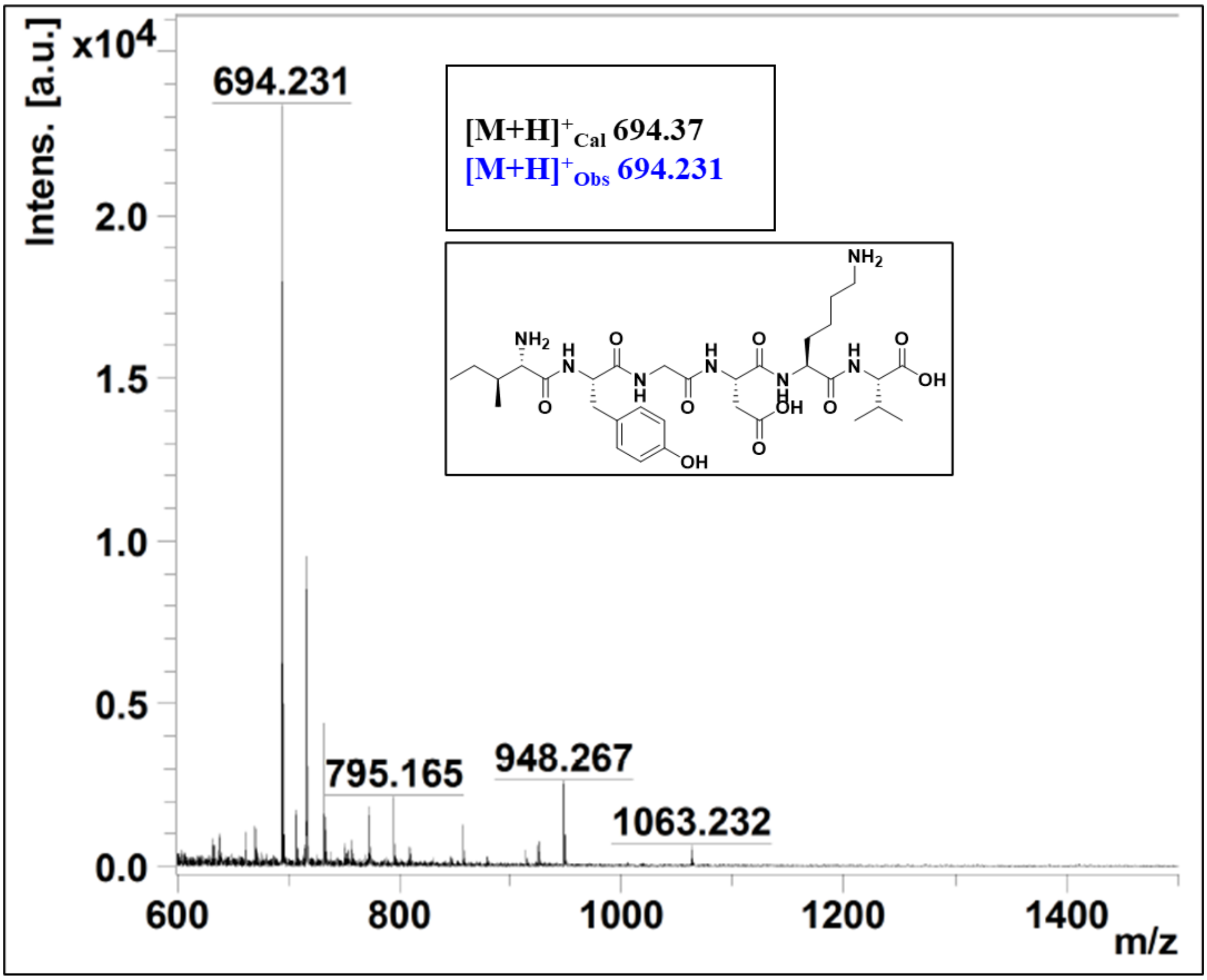


**Figure S13.** MALDI spectrum of H-Ile-Tyr-Gly-Asp-Lys-Val-OH in a mixture of acetonitrile: water, expected molecular weight is 694.37, and the observed molecular weight *via* MALDI is 694.231.

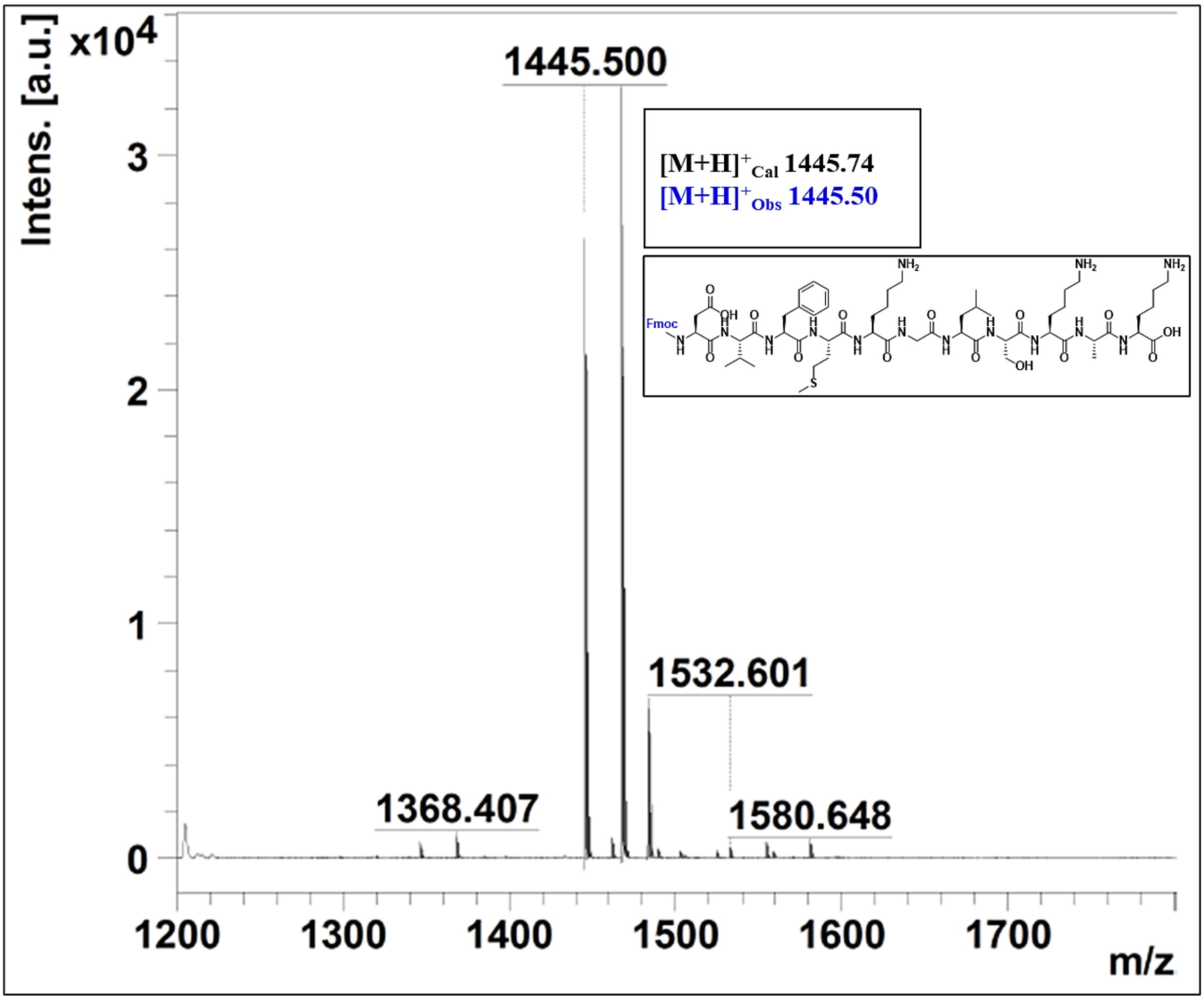


**Figure S14.** MALDI spectrum of Fmoc-Asp-Val-Phe-Met-Lys-Gly-Leu-Ser-Lys-Ala-Lys-OH in a mixture of acetonitrile: water, expected molecular weight is 1445.74, and the observed molecular weight *via* MALDI is 1445.500.

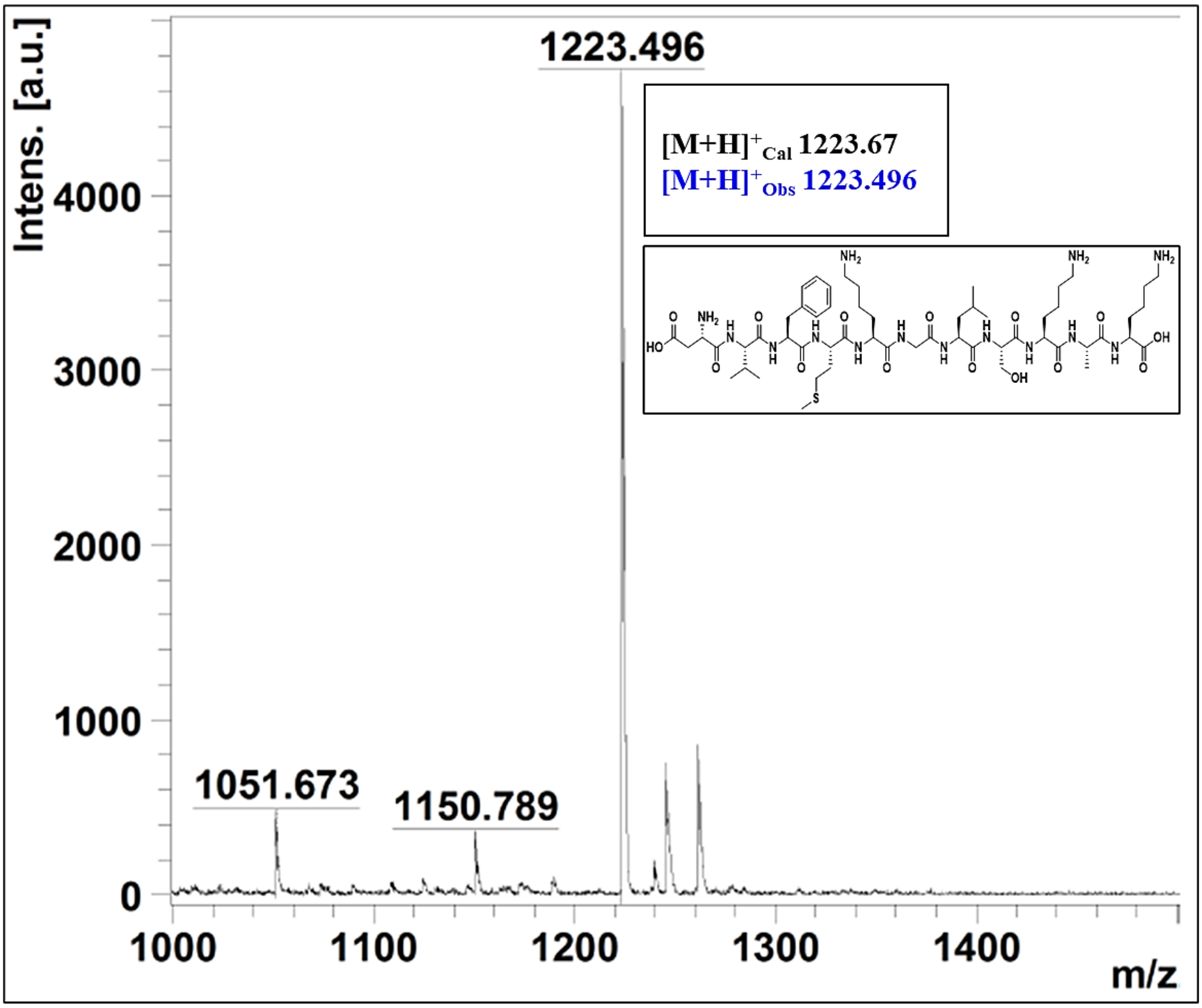


**Figure S15.** MALDI spectrum of H-Asp-Val-Phe-Met-Lys-Gly-Leu-Ser-Lys-Ala-Lys-OH in a mixture of acetonitrile: water, expected molecular weight is 1222.67, and the observed molecular weight *via* MALDI is 1223.496.

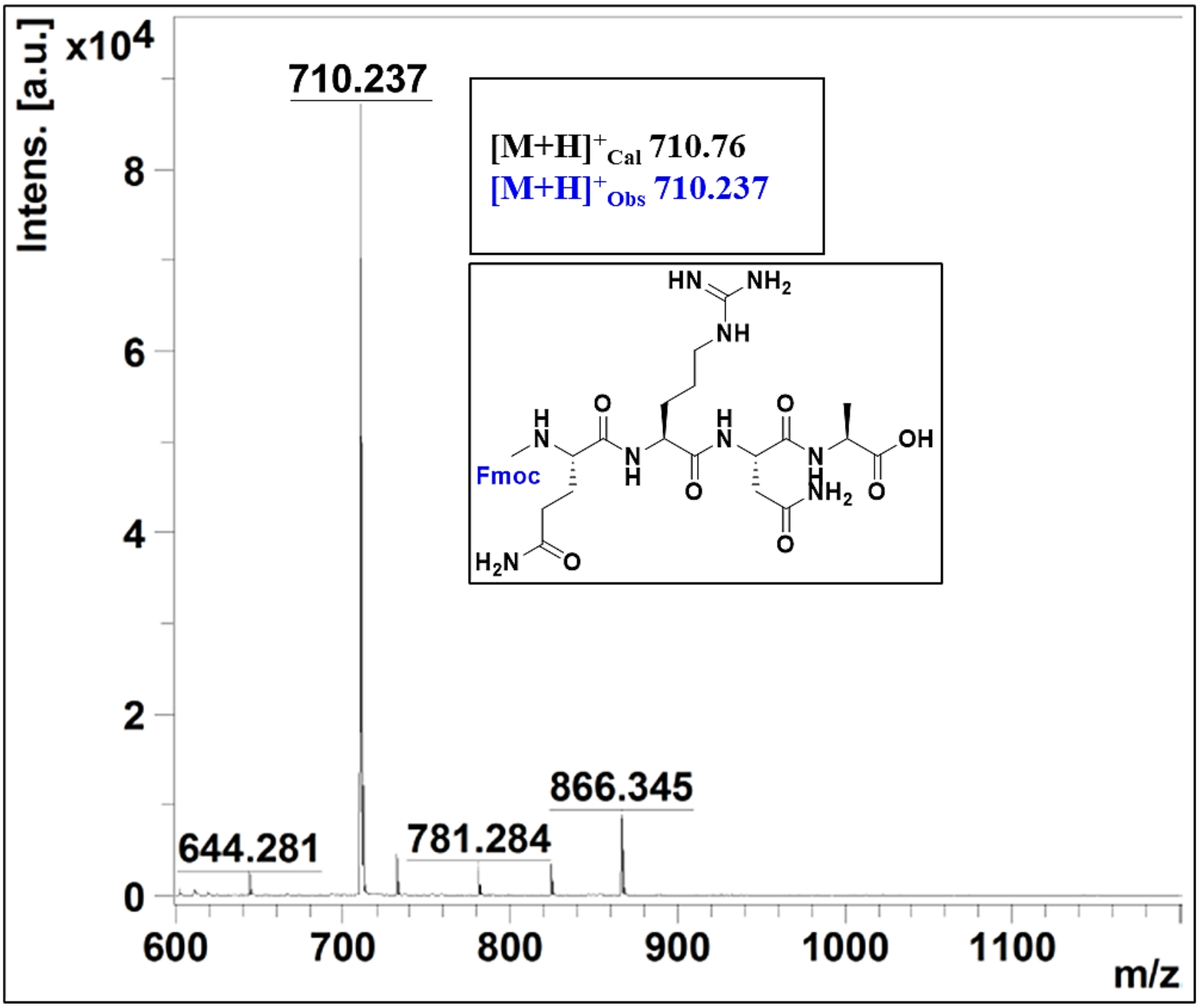


**Figure S16.** MALDI spectrum of Fmoc-Gln-Arg-Asn-Ala-OH in a mixture of acetonitrile: water, expected molecular weight is 710.76, and the observed molecular weight *via* MALDI is 710.237.

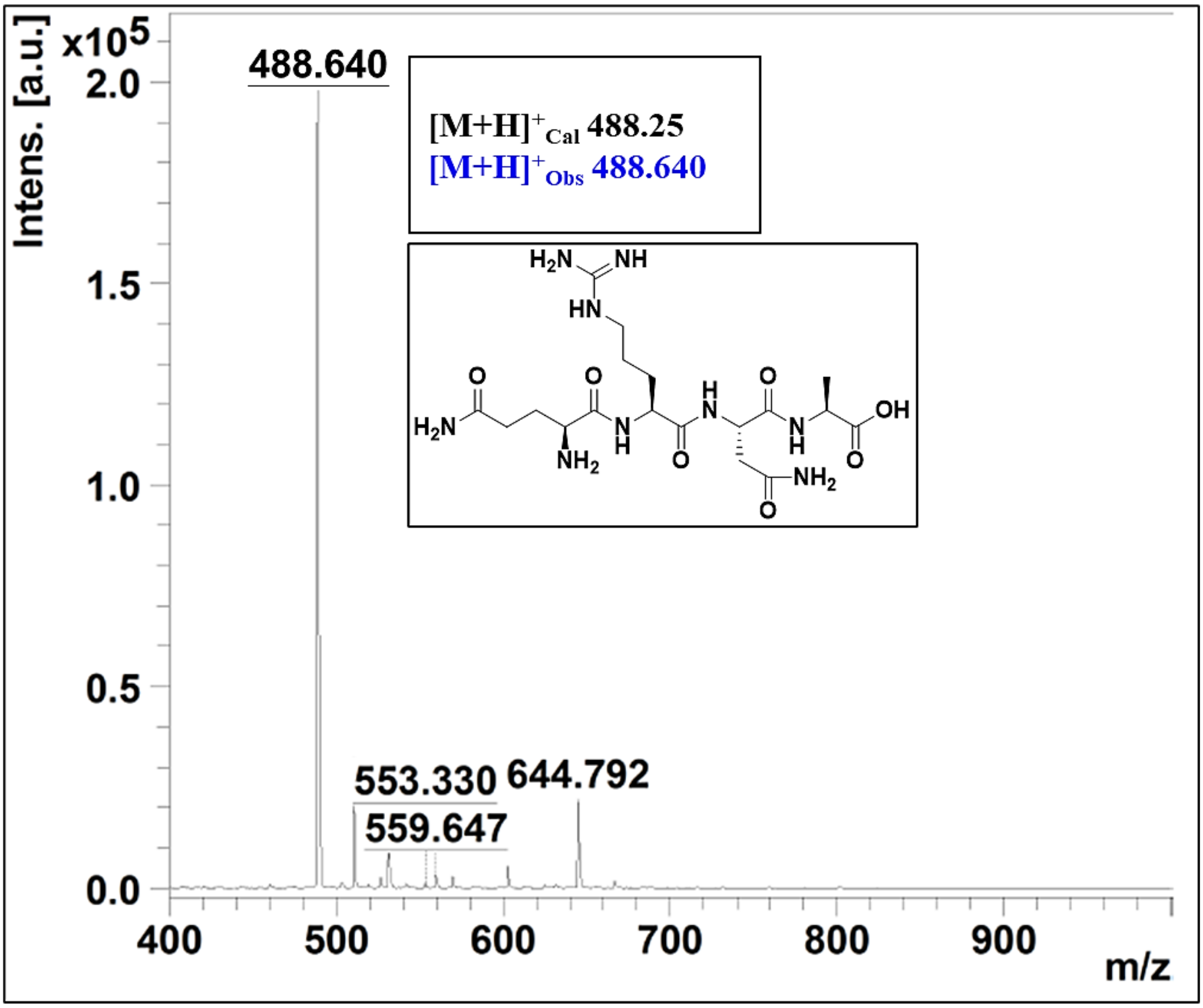


**Figure S17.** MALDI spectrum of H-Gln-Arg-Asn-Ala-OH in a mixture of acetonitrile: water, expected molecular weight is 488.25, and the observed molecular weight *via* MALDI is 488.640.

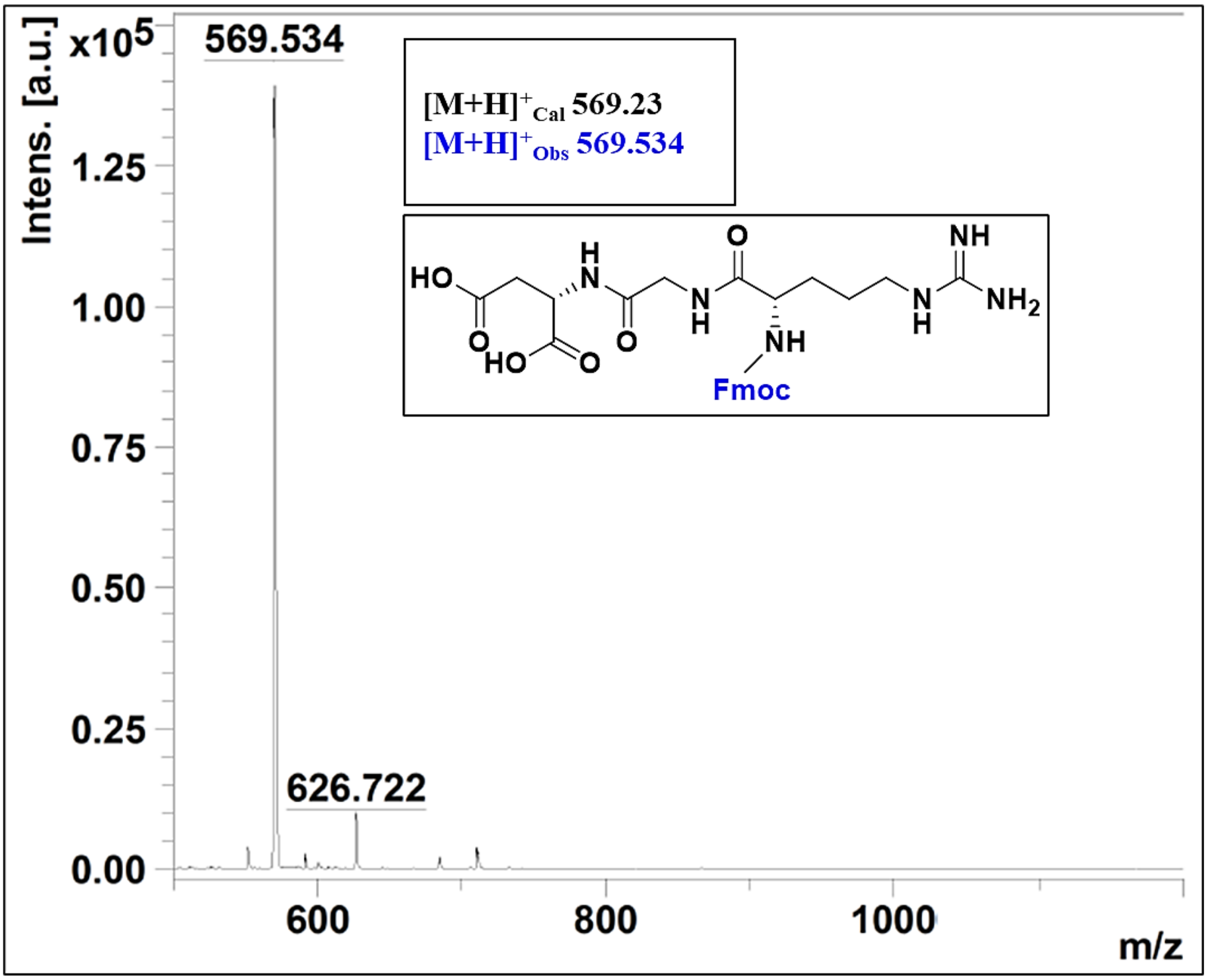


**Figure S18.** MALDI spectrum of Fmoc-Arg-Gly-Asp-OH in a mixture of acetonitrile: water, expected molecular weight is 569.23, and the observed molecular weight *via* MALDI is 569.534.

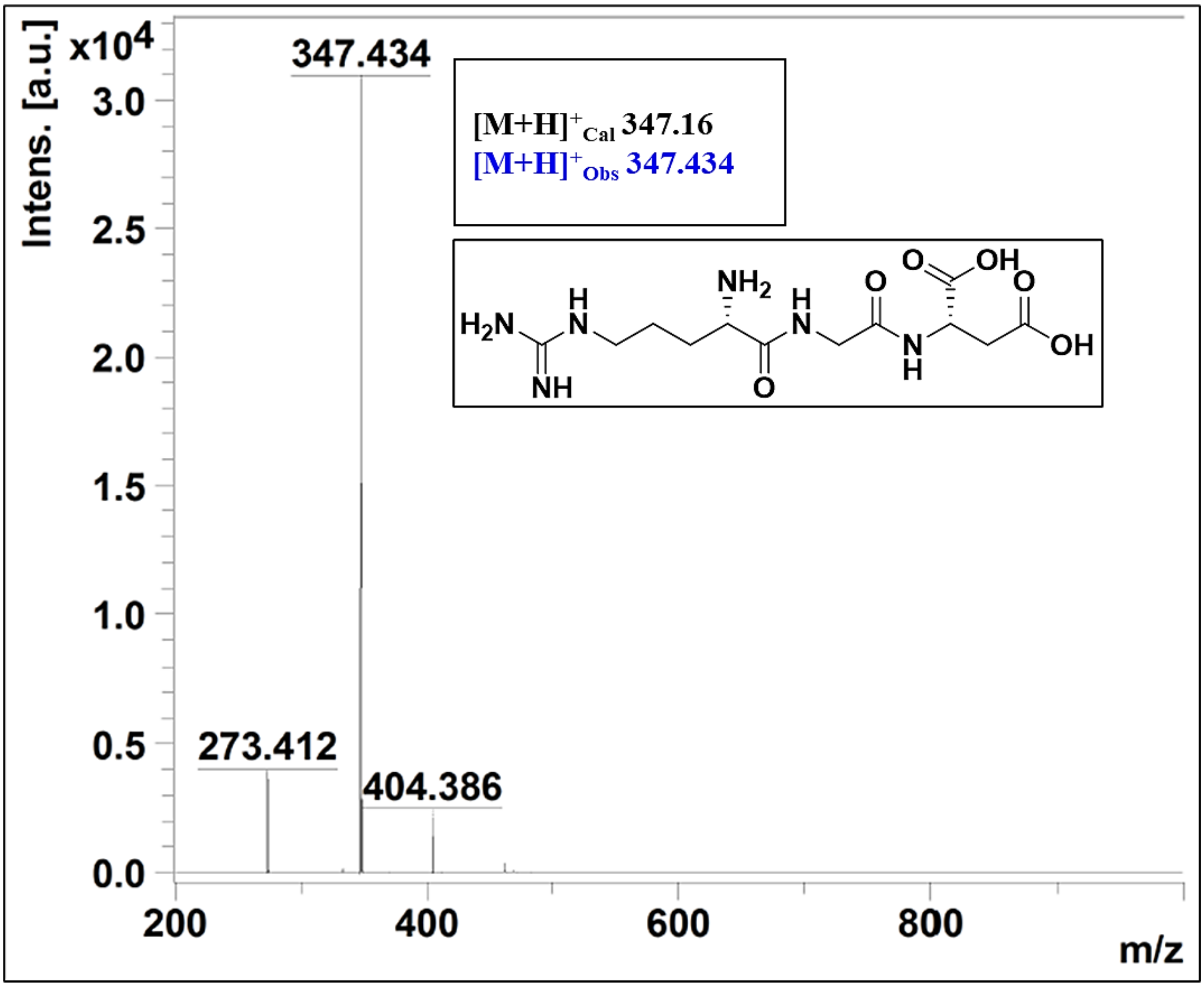


**Figure S19.** MALDI spectrum of H-Arg-Gly-Asp-OH in a mixture of acetonitrile: water, expected molecular weight is 347.16, and the observed molecular weight *via* MALDI is 347.434.
